# Multimodal 3D imaging reveals a central role for lysosomes in dissolution of cholesterol crystals by macrophages

**DOI:** 10.64898/2026.09.22.753145

**Authors:** Vibeke Akkerman, Alice Dupont Juhl, Jacob M. Egebjerg, Christoph Pratsch, Stephan Werner, Gerd Schneider, Tido Willms, Peter Müller, Sergey Kapishnikov, Daniel Wüstner

**Affiliations:** Department of Biochemistry and Molecular Biology, University of Southern Denmark, Campusvej 55, DK–5230 Odense M, Denmark; Department of X-Ray Microscopy, Helmholtz-Zentrum Berlin, Albert-Einstein-Str. 15, 12489 Berlin, Germany; Department of Biology, Humboldt University Berlin, Invalidenstr. 42, 10115 Berlin, Germany; SiriusXT Limited, Dublin, Ireland

**Keywords:** cholesterol crystals, digestive exophagy, macrophages, cyclodextrin, fluorescence, atherosclerosis, microscopy, soft X-ray microscopy

## Abstract

Formation of cholesterol crystals (CCs) is a key event during the development of atherosclerosis, but the molecular mechanisms of their degradation within cells are poorly understood. By incorporating the fluorescent cholesterol analogue TopFluor-Cholesterol (TF-Chol) into CCs, we were able to visualize the uptake of CCs in macrophages using correlative fluorescence and soft X-ray microscopy. Using quantitative 3D live-cell imaging, we show that CCs are processed in late endosomes and lysosomes (LE/Lys), resulting in formation of TF-Chol containing lipid droplets (LDs) over time. Inhibition of lysosomal sterol export with U18666A caused accumulation of TF-Chol in LE/Lys, and inhibition of lysosomal acidification with bafilomycin A1 led to reduced dissolution of the CCs. Using a novel assay combined with 3D image processing, we show that large CCs in contact with macrophages are processed via lysosomal exocytosis followed by extracellular and intracellular degradation of CCs. Treating macrophages with a fluorescent version of cyclodextrin (CD) promoted the dissolution of CCs and enhanced the formation of LDs enriched with TF-Chol. The majority of fluorescent CD co-localized with a marker for LE/Lys during this process, suggesting that intracellular delivery to LE/Lys may contribute to the dissolution of CCs. Dehydroergosterol (DHE) is an intrinsically fluorescent sterol closely mimicking the properties and behavior of cholesterol. DHE is known to self-associate into aggregates and crystals, and by using fluorescence spectroscopy and specialized ultraviolet (UV) microscopy, we found that CD enhances the dissolution of DHE crystals *in vitro* and in cells. Together, our findings highlight the lysosomal pathway as responsible for dissolution of CCs, cholesterol trafficking, and efflux in macrophages.

**For Table of contents only:** 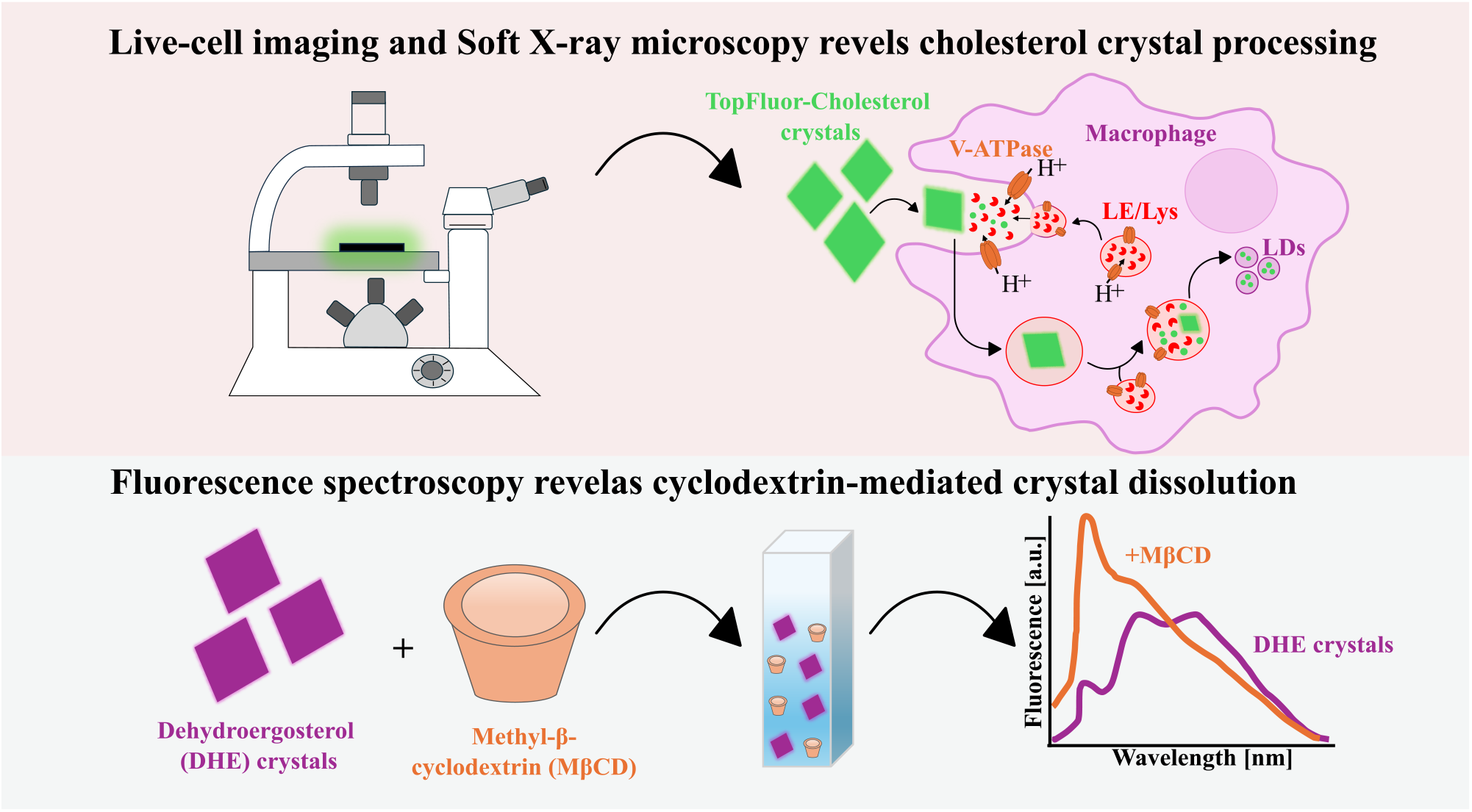

## Introduction

Atherosclerosis is a chronic inflammatory disease associated with elevated cholesterol levels, and it is the pathology that causes cardiovascular diseases, including heart attack and stroke, one of the leading causes of death worldwide.^1^ The disease affects the arteries, especially the intima layer, where circulating low-density lipoproteins (LDL) are deposited, modified, and aggregated, activating monocytes that differentiate into macrophages.^2,3^ Macrophages process aggregated LDL (agLDL) by forming deep invaginations in which they secrete hydrolytic enzymes from lysosomes, a process called lysosomal exocytosis. This results in the hydrolysis of cholesteryl esters (CEs) within agLDL and the generation of large amounts of unesterified (free) cholesterol. High cholesterol uptake by macrophages stimulates intracellular cholesterol re-esterification by acyl-CoA:cholesterol acyltransferase (ACAT), and the resulting CEs are stored in lipid droplets (LDs). This excessive amount of CEs leads to the conversion of macrophages into foam cells, a critical step and hallmark of early atherosclerosis.^2–4^ Due to the limited solubility of free cholesterol in lipid membranes and in lipoproteins, excessive cholesterol can precipitate, and cholesterol crystals (CCs) can form both extracellularly and intracellularly.^5^ Extracellular crystals arise from lipoprotein aggregation within the intima, where hydrolysis of cholesteryl esters in retained and aggregated LDL generates high concentrations of unesterified cholesterol,^4^ whereas intracellular crystals may develop during lysosomal degradation of modified lipoproteins or when the capacity of ACAT-mediated cholesterol esterification is exceeded.^6,7^ CCs were traditionally considered a feature of advanced atherosclerotic lesions; however, smaller CCs have also been identified in early atherosclerotic lesions.^5,8^ Furthermore, the dissolution rate of CCs in patient serum has been shown to predict the outcome of surgical intervention for aortic stenosis.^9^ Cellular uptake of CCs triggers a complex network of inflammatory responses, including the activation of NLRP3 inflammasomes, thereby promoting cytokine secretion and further progression of atherosclerosis.^10,11^ CCs can adopt a variety of morphologies, some of which can eventually perforate the endothelial layer, resulting in thrombus formation, a hallmark of advanced atherosclerosis^5,10,12,13^.

Because CCs contribute to plaque progression and inflammation, strategies that promote their dissolution have attracted considerable interest. Cyclodextrins (CDs) are cyclic oligosaccharides with a hydrophilic surface and a hydrophobic central cavity known to form inclusion complexes with cholesterol.^14^ There are three major classes of CDs defined by the number of sugar units in the ring structure. *β*-cyclodextrins, composed of seven sugar units, are widely used to increase the bioavailability of poorly water-soluble drugs and sterol compounds.^14,15^ Studies in murine models of atherosclerosis have shown that CD treatment reduces atherosclerotic plaque size.^16^ The mechanisms, by which CDs act, remain debated. CDs are known to extract cholesterol from the plasma membrane (PM) of cultured cells, and one hypothesis is that intracellular cholesterol-rich lysosome-like organelles transport cholesterol to the PM, which then can be further removed by CDs.^17^ Another hypothesis is that CDs are internalized by cells through endocytosis or pinocytosis and delivered to the late endosomes/lysosomes (LE/Lys), from where cholesterol can be transported to other organelles.^18^ A third possibility is, that CDs trigger lysosomal exocytosis and thereby secretion of cholesterol-rich membranes.^19,20^ The mechanisms by which CDs promote intracellular CC dissolution remain poorly understood.

Currently, few tools are available for investigating the cellular mechanisms underlying uptake and degradation of CCs. Dehydroergosterol (DHE), an intrinsically fluorescent analog of cholesterol, can form sterol-rich aggregates and crystalline structures.^21^ This analog is minimally modified compared to cholesterol, with only two additional double bonds in the ring structure, and an extra methyl group and doule bond in the side chain. However, DHE is fluorescent in the ultraviolet (UV) range of the spectrum, resulting in its visualization requiring specialized UV-sensitive fluorescence microscopes.^22^ TopFluor-cholesterol (TF-Chol) is another fluorescent analog of cholesterol, which has a boron-dipyrromethene (BODIPY) moiety linked to carbon-24 of the sterol side chain. It emits green fluorescence and is more photostable compared to DHE, enabling its visualization using conventional fluorescence microscopy systems.^23^ However, the attachment of a fluorescent group may alter the properties of the analog compared with native cholesterol.^22–24^

In this study, we combine imaging of both TF-Chol and DHE in sterol crystals to map the pathways underlying processing of CCs in macrophages. Preliminary evidence suggests that this combined approach can shed new light onto processing of CCs in macrophages.^25^ Here, we demonstrate that doping CCs with TF-Chol enables 3D confocal imaging and soft X-ray microscopy of CC uptake and dissolution in cells. By adapting a lysosomal exocytosis assay using biotin-conjugated fluorescent dextran and TF-Chol-doped CCs, we show that lysosomal contents are delivered to the surface of larger CCs, as previously observed with agLDL.^26–28^ This process of digestive exophagy of extracellular CCs was reduced in the presence of bafilomycin A1 (Baf-A1), an inhibitor of vacuolar-type *H*^+^ (V-ATPase), demonstrating the requirement of lysosomal acidification for efficient lysosomal exocytosis. Furthermore, we identify LE/Lys as key compartments involved in CC processing and dissolution. Using fluorescent methyl-*β*-cyclodextrin (M*β*CD), we demonstrate that cholesterol extraction from CCs occurs within LE/Lys in macrophages, promoting the formation of TF-Chol-enriched LDs. Inhibition of lysosomal cholesterol transport and degradation pathways further highlights the essential role of LE/Lys in CC processing. Finally, using DHE as a minimally modified fluorescent cholesterol analog, we confirmed that M*β*CD treatment promotes sterol crystal dissolution *in vitro* and in living macrophages.

## Results

### Uptake and Digestion of Cholesterol Crystals by Macrophage Cells

Macrophages are phagocytic cells of the innate immune system and play a key role in the development of atherosclerosis.^29^ Here, we used the transformed murine macrophage cell line, J774, which is widely used in studies of atherosclerosis.^3,26^ To investigate uptake and processing of CCs, J774 macrophages were incubated with CCs doped with TF-Chol (TF CCs) for either 5 or 16 h before live-cell 3D confocal imaging. After 5 h, we observed partial crystal degradation in macrophages, and in cells associated with crystals, TF-Chol fluorescence localized to both LE/Lys and LDs (Figure 1). To quantify these observations, we developed a 3D image analysis workflow incorporating a machine-learning model trained to segment LDs, enabling quantitative analysis of multi-color confocal image stacks (see Materials and Methods for details). By measuring the fraction of the cell volume occupied by LDs and LE/Lys after 5 and 16 h of incubation with TF CCs, we found that overnight incubation resulted in a lower volume fraction of both organelles compared with 5 h exposure to TF CCs. However, this reduction in organelle volume fraction may partly reflect the prolonged exposure to lipoprotein-deficient serum (LPDS) medium rather than crystal processing alone. Additionally, the mean TF-Chol fluorescence intensity in both LDs (89.8 ± 14.2 vs. 67.3 ± 8.0) and LE/Lys (75.2 ± 11.4 vs. 41.3 ± 4.4) was higher after 5 h than after overnight incubation (Figure S1), suggesting progressive TF-Chol efflux over time.

**Figure 1:**
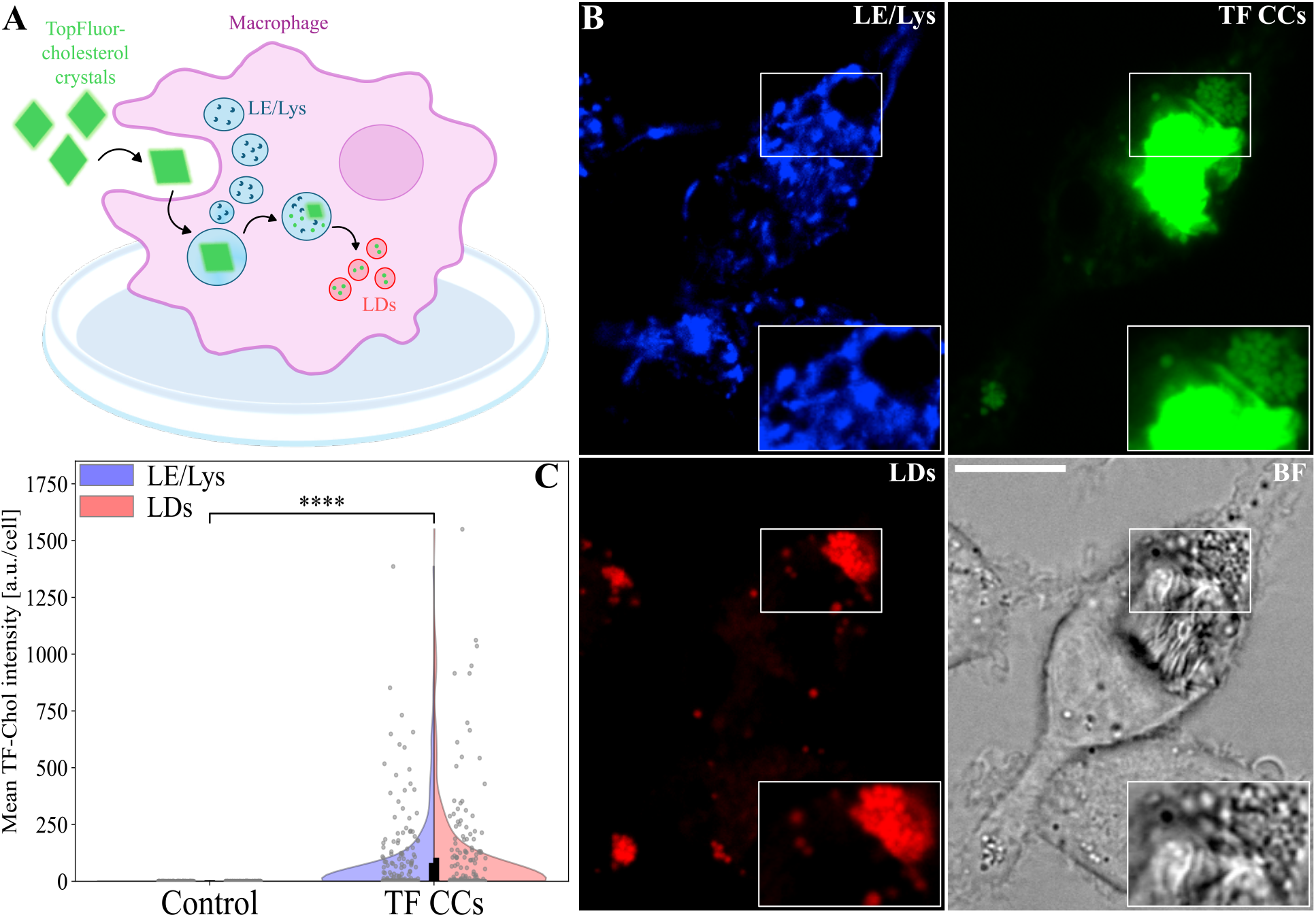
Uptake and intracellular trafficking of TopFluor-Cholesterol crystals (TF-CCs) in J774 macrophages. Schematic illustration of macrophages treated with TF-CCs, and the intracellular trafficking to late endosomes/lysosomes (LE/Lys) and lipid droplets (LDs) (A). Representative live-cell confocal microscopy images of macrophages incubated with TF CCs for 5 h in DMEM/LPDS medium. LE/Lys compartments were labeled with Cascade Blue (blue), and LDs were stained with LipidTOX (red). TF-Chol fluorescence is shown in green (TF-CCs). Scale bar, 10 µm (B). Quantification of TF-Chol fluorescence intensity associated with LE/Lys (blue) and LDs (red) per cell in untreated control macrophages and TF-CC-treated cells. Data represent mean fluorescence intensity per cell from two independent experiments for control cells (n = 141 cells) and three independent experiments for TF-CC-treated cells (n = 219 cells). Statistical significance was assessed using the Mann-Whitney U test: control vs. TF-CCs, p = 5.08 *×* 10*^−^*^55^ (LE/Lys), and 1.59 *×* 10*^−^*^45^ (LDs).

Time-lapse live-cell imaging of macrophages in the presence of TF CCs overnight further revealed that the macrophages exhibit cooperative behavior in order to digest CCs (Figure S2). The macrophages were co-stained with the LE/Lys marker, Rh-dextran, and we observed that the LE/Lys clustered around the crystals from the outset (Figure S2, white arrows), consistent with a role for these compartments in crystal processing. We generally observed a close association between LE/Lys and TF CCs (Movie 1). Crystal size is likely to influence the uptake and degradation mechanism. Smaller crystals are most likely internalized by endocytosis, whereas processing of larger crystals could also involve a form of extracellular digestion.

Confocal microscopy does not have sufficient resolution to resolve the ultrastructure of small CCs nor the exact mechanism of uptake of small and large CCs. Since TF-CCs contain mostly cholesterol and only small amounts of the fluorescent sterol, they can be visualized by correlative fluorescence and soft X-ray microscopy. The latter technique is also called soft Xray tomography (SXT), and its contrast is based on the absorption of X-rays by carbon-rich materials, such as CCs. Due to the use of much shorter wavelengths than used in fluorescence microscopy, SXT has a much higher resolution of about 40 nm in all three directions.^30^ This makes SXT ideally suited to study the ultrastructure of internalized CCs (Figure 2 and Figure S3). SXT carried out at a lab-based X-ray microscope revealed elongated crystal structures within macrophages,^31^ and the corresponding fluorescence signal confirmed that these structures contained TF-Chol (Figure 2A-C, F-H). A montage of consecutive slices of the tomogram reconstruction further demonstrated that a larger crystal extended through the cellular volume, was partially ingested and surrounded by cellular material (Figure 2D). Similar crystal-containing macrophages were observed in synchrotron-based SXT datasets acquired at BESSY II, where correlative TF-Chol fluorescence and X-ray imaged from a reconstructed 3D volume confirmed the intracellular localization of smaller crystals (Figure 2F-H and). Interestingly, such rod-shaped CCs often appeared aggregated and close to the nucleus (Figure 2E-H and S3). Together, these results confirm that macrophages internalize small CCs and transport them towards the cell center, but engulf large CCs only partially.

**Figure 2:**
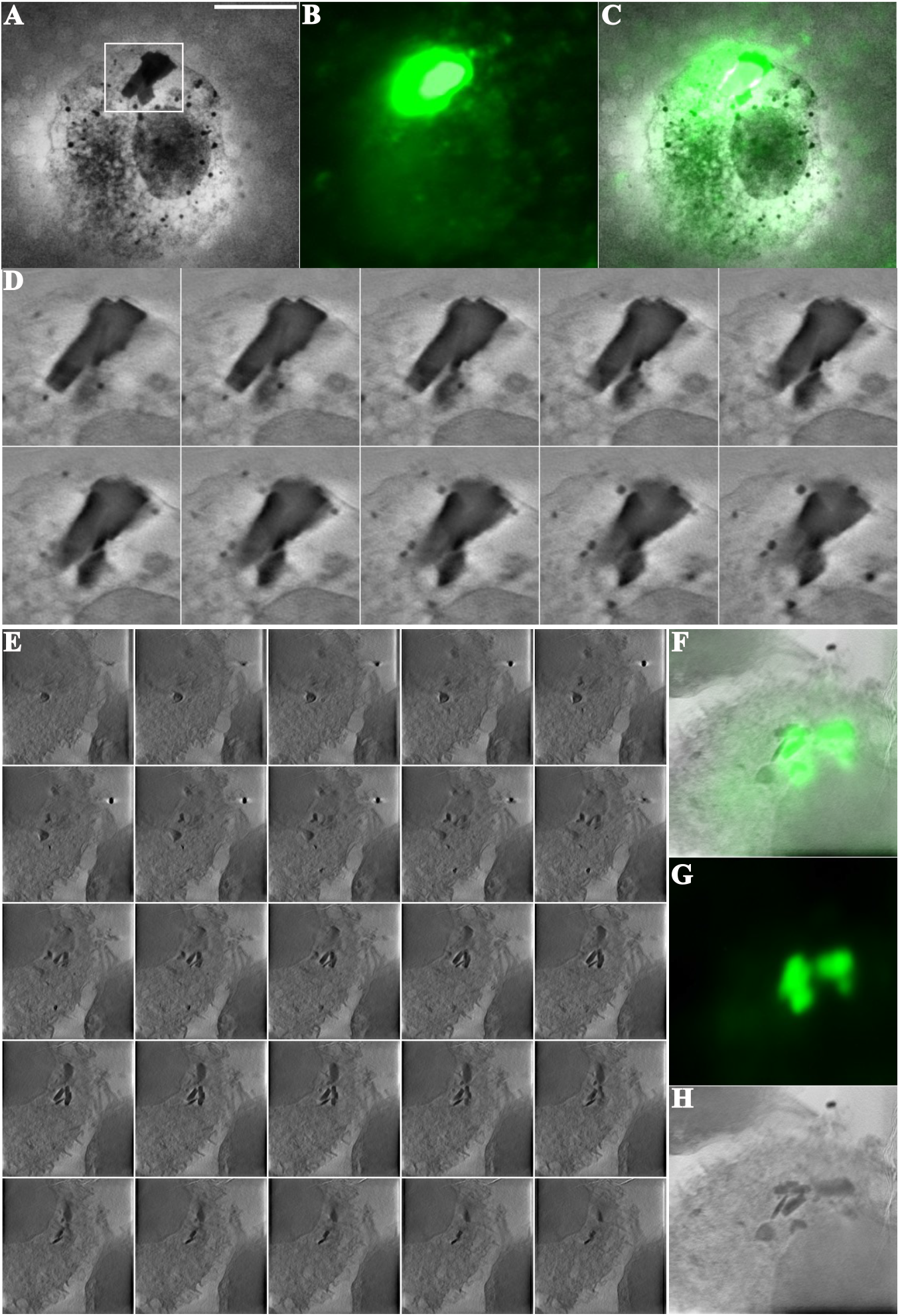
Correlative fluorescence and soft X-ray of crystal uptake and processing by macrophages. J774 macrophages were seeded onto grids and cultured for 2 days before overnight incubation with TF CCs. Representative macrophage containing an internalized TF CC imaged using the Sirius XT SXT100, shown as an SXT volume slice (A), epifluorescence image (B), and overlay (C). Scale bar, 10 µm. (D) Higher-magnification views of the crystal in (A-C) shown in consecutive volume slices. SXT of TF CCs ingested by macrophages was also carried out at the BESSY II synchrotron at Helmholtz Zentrum Berlin (HZB) (E-H). Representative tomographic slices through the 3D reconstruction (E) and correlative images of TF-Chol fluorescence in crystals together with the corresponding SXT sum projection of the same depth of field (F-H).

### Disruption of Lysosomal Function Impairs Cholesterol Crystal Processing and Promotes Intracellular TF-Chol Accumulation

The LE/Lys are central regulators of intracellular cholesterol homeostasis. Cholesterol, obtained for example from hydrolysis of lipoprotein-derived cholesteryl esters, is exported from LE/Lys primarily through the coordinated actions of NPC2 and the transmembrane transporter NPC1.^32,33^ This process depends also on acidification of the lysosomal lumen by the vacuolar-type *H*^+^-ATPase (V-ATPase). To investigate whether lysosomal function is required for cholesterol crystal processing, macrophages were treated with bafilomycin A1 (Baf-A1), a macrolide antibiotic that inhibits the V-ATPase, thereby preventing lysosomal acidification and impairing lysosomal degradation. Cells were treated with Baf-A1 for 1 h before treatment with CCs for 5 h (Figure 3). We found that Baf-A1-treated cells were still able to internalize CCs, indicating that Baf-A1 does not prevent CC uptake, consistent with a previous report.^34^ However, TF-Chol fluorescence was markedly reduced in cellular compartments such as LDs and LE/Lys following Baf-A1 treatment (Figure 3). No significant difference was observed in the fraction of cell volume occupied by LDs or LE/Lys between control and Baf-A1-treated cells (data not shown). Specifically, mean TF-Chol fluorescence intensity in LE/Lys decreased from 75.2 ± 11.4 to 25.7 ± 4.7, while TF-Chol fluorescence in the LDs decreased from 89.8 ± 14.2 to 23.2 ± 4.8 (Figure 3C). In addition, we determined the fraction of total cellular TF-Chol fluorescence detected within LE/Lys or LDs, calculated as the TF-Chol fluorescence within the respective organelle mask relative to the total TF-Chol fluorescence within the cell. The fraction was significantly reduced for LE/Lys following BafA1 treatment, whereas the fraction detected within LDs remained unchanged (Figure 3D). Together, these findings suggest that lysosomal acidification is required for efficient TF CC processing and regulates the intracellular redistribution of TF-Chol into intracellular cholesterol pools. This is consistent with previous reports showing that Baf-A1 inhibits lysosomal degradation, resulting in the persistence of undigested and partly phagocytosed material.^34^

**Figure 3:**
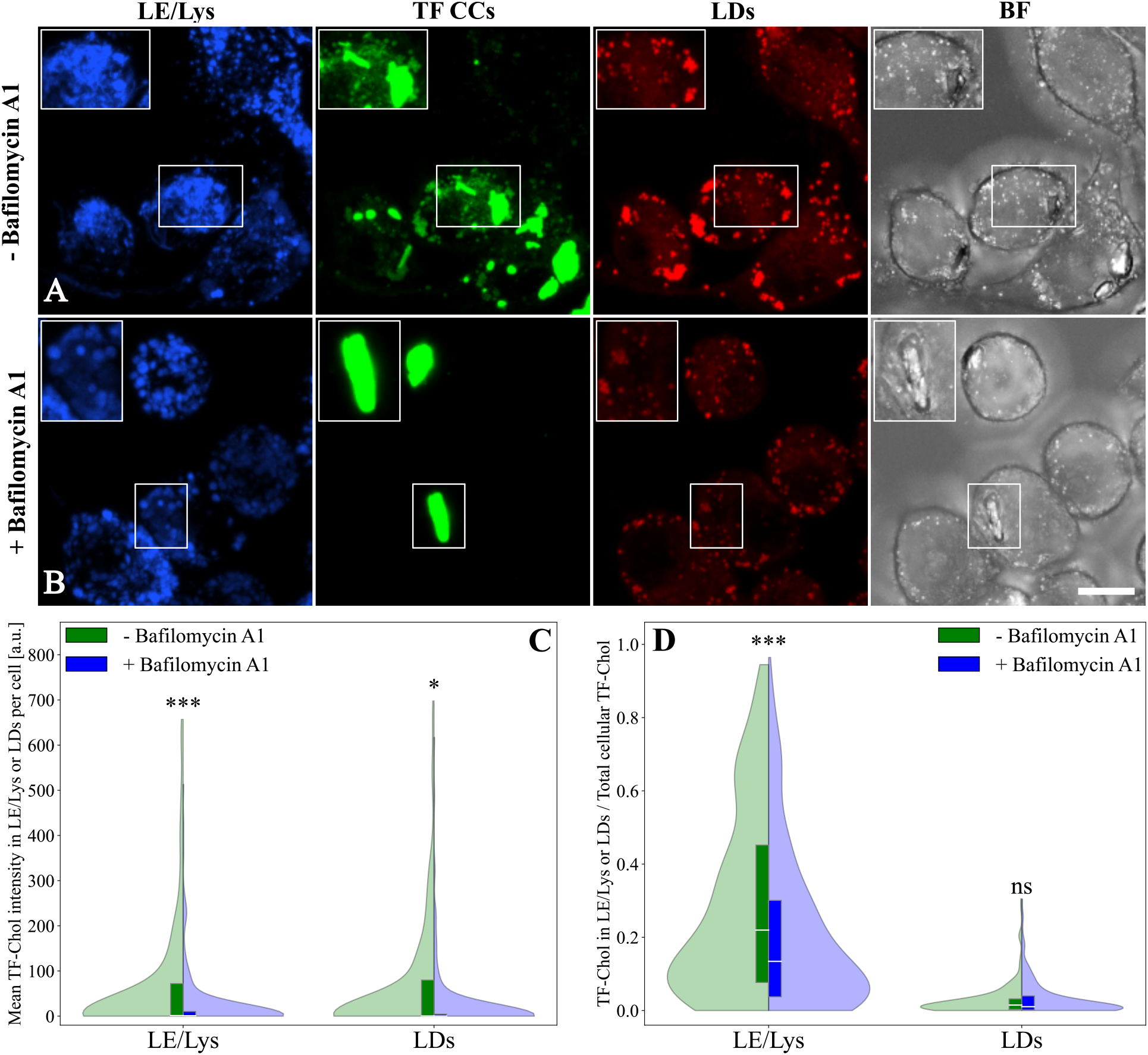
Lysosomal processing of cholesterol crystals is impaired by bafilomycin A1 (Baf-A1), a V-ATPase inhibitor. Representative confocal maximum intensity projection images of macrophages treated with Baf-A1 1 h before exposure to TF CCs for 5 h (A-B). Scale bar, 10 µm. Mean TF-Chol fluorescence intensity in LE/Lys and LDs per cell in untreated (green) and Baf-A1-treated cells (blue) (C). Fraction of total cellular TF-Chol fluorescence localized within LE/Lys and LDs (D). Statistical significance was determined using Mann-Whitney U test: p = 3.83 *×* 10*^−^*^4^ (C, LE/Lys), p = 3.92 *×* 10*^−^*^2^ (C, LDs), p = 3.29 *×* 10*^−^*^4^ (D, LE/Lys), and p = 5.82 *×* 10*^−^*^1^ (D, LDs). Quantification was performed for three independent experiments for untreated cells (n = 219 cells) and Baf-A1-treated cells (n = 222 cells).

Since lysosomal acidification was required for efficient crystal processing, we next investigated whether cholesterol export from LE/Lys contributes to TF-Chol redistribution. To determine whether cholesterol crystal-derived TF-Chol exits LE/Lys through NPC1-dependent transport, J774 macrophages were incubated overnight with TF CCs in the presence of U18666A, an inhibitor of NPC1-mediated cholesterol export from the LE/Lys. U18666A resulted in enlarged LE/Lys (Figure S4) (0.037 ± 0.0016) compared with TF CC-treated cells (0.028 ± 0.0009). In addition, TF-Chol fluorescence within the LE/Lys was markedly increased following U18666A treatment (126.1 ± 12.8) relative to TF CC treatment alone (41.3 ± 4.4), indicating that TF-Chol accumulates in the LE/Lys when NPC1-mediated cholesterol export is inhibited. This is consistent with previous studies demonstrating that U18666A does not affect LDL binding, internalization, or lysosomal hydrolysis of LDL-cholesteryl esters, but instead causes lysosomal accumulation of cholesterol by preventing its transport to other cellular membranes.^35^ Interestingly, we found that the LD volume fraction was significantly increased in U18666A-treated cells (0.0120 ± 0.0005) compared with untreated cells (0.0046 ± 0.003) (Figure S4C). However, despite the increased LD volume fraction, TF-Chol fluorescence within LDs showed only a minor increase (Figure S4D). These findings indicate that U18666A-induced LD volume fraction is not accompanied by proportional TF-Chol accumulation, suggesting that LD expansion may represent a broader adaptive response to altered cholesterol homeostasis rather than simply reflecting direct accumulation of crystal-derived cholesterol.

### Processing of Large Cholesterol Crystals by Macrophages Involves Lysosomal Exocytosis

Digestive exophagy is a process by which macrophages degrade extracellular objects that are too large to be internalized by conventional phagocytosis. During this process, macrophages generate an extracellular acidic, hydrolytic compartment through lysosomal exocytosis to facilitate degradation of large extracellular substrates, such as agLDL. A previously established assay using lysosome-loaded biotin-dextran and streptavidin-labelled agLDL enables detection of lysosomal content delivery through biotin-streptavidin-mediated capture of released lysosomal material.^26–28^

The size of experimentally generated CCs is heterogeneous, and crystal size is likely to influence the mechanism by which macrophages process them. To investigate whether larger CCs can be processed by macrophages through digestive exophagy, we developed a strategy to detect lysosomal exocytosis at the crystal surface. Because cholesterol crystals have smooth, chemically inert surfaces lacking functional groups suitable for covalent modification, such as amine, carboxyl, or thiol groups, we incorporated cholesterol-PEG-biotin into the crystal structure. The PEG spacer exposes the biotin moiety at the crystal surface, enabling NeutrAvidin binding while leaving available biotin-binding sites for capture of exocytosed biotin-labelled dextran. This strategy enables detection of lysosomal content delivery to the cholesterol crystal surface during exophagy. Following the strategy previously established for agLDL, J774 macrophages were incubated overnight with 1 mg/ml biotin-rhodamine-dextran, followed by a 3 h chase period to allow delivery to lysosomes. Baf-A1 was added 1 h prior to exposure to NeutrAvidin-biotin-TF-CCs. Cells were subsequently incubated with excess biotin in medium to bind any unoccupied NeutrAvidin sites and fixed before imaging on a confocal microscope (Figure 4).

**Figure 4:**
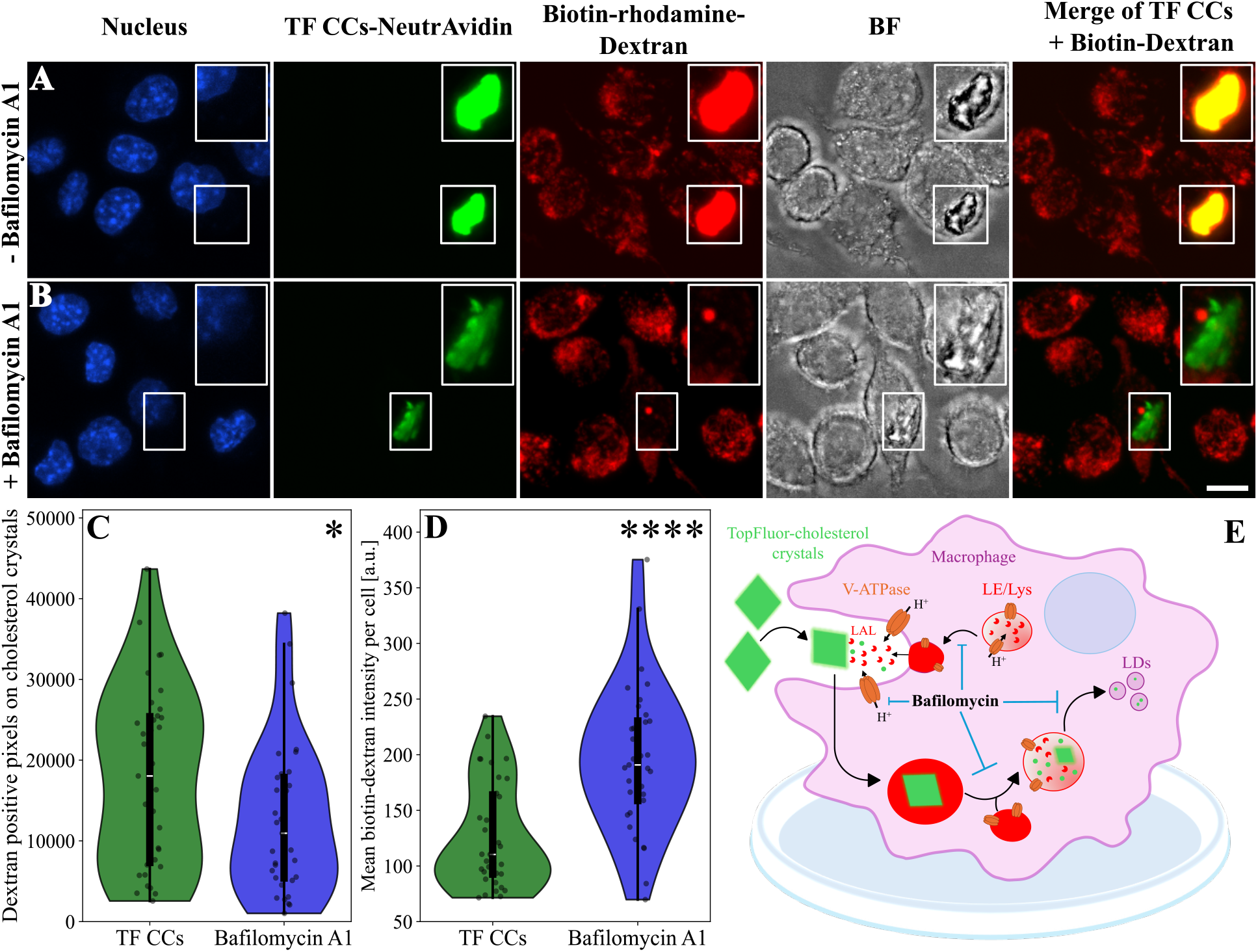
Macrophages deliver lysosomal contents to cholesterol crystals through digestive exophagy. J774 macrophages were incubated overnight with 1 mg/ml biotin-rhodamine-dextran, followed by a 3 h chase to allow delivery of biotin-dextran to lysosomes. Baf-A1 was added 1 h before exposure to biotin-functionalized TF-Chol cholesterol crystals (TF CCs) coated with NeutrAvidin for 90 min. Biotin-dextran released by lysosomal exocytosis is captured at the crystal surface through the remaining biotin-binding sites of NeutrAvidin. Representative maximum-intensity projection of macrophages exposed to TF CCs (A) or pre-treated with Baf-A1 before TF CCs exposure (B). Scale bar, 10 µm. Quantification was performed from three independent experiments. The area of biotin-dextran-positive signal overlapping with cholesterol crystals (CCs) was quantified (C, p = 2.99 *×* 10*^−^*^2^). Mean intracellular biotin-dextran intensity per cell was quantified (D, p = 1.18 *×* 10*^−^*^5^). Schematic illustration of the proposed effect of Baf-A1 (E).

After 90 min of incubation with crystals, biotin-dextran fluorescence was observed at the crystal surface (Figure 4). This was further supported by quantification of the area of crystal-associated biotin-dextran fluorescence, which was significantly increased in CCs-treated cells (18020.06 ± 1868.3) compared with control cells without dextran (972.3 ± 294.4), used to assess potential signal crosstalk (data not shown). In contrast, Baf-A1-treated cells showed reduced accumulation of biotin-dextran on crystals (12603.9 ± 1630.3) (Figure 4C), indicating reduced delivery of lysosomal contents to the crystal surface. Interestingly, the mean intracellular dextran intensity per cell was increased in Baf-A1-treated cells (196.3 ± 11.3) compared with cells exposed to crystals alone (126.0 ± 7.9) (Figure 4D). Together, these findings indicate that Baf-A1 reduces the delivery of lysosomal contents to cholesterol crystals, as reflected by decreased accumulation of biotin-dextran at the crystal surface and increased intracellular biotin-dextran fluorescence.

### Cyclodextrin Facilitates the Dissolution of Cholesterol Crystals upon Uptake into Endo-lysosomes

Having established that macrophages process cholesterol crystals through lysosomal pathways, we next investigated whether cyclodextrin promotes intracellular crystal dissolution. To this end, macrophages were incubated with TF CCs for 5 h to allow crystal uptake before treating cells overnight with a mixture of fluorescent and non-fluorescent M*β*CD (Figure 5). Following M*β*CD treatment, most intracellular CCs were no longer detectable, and TF-Chol fluorescence was redistributed from crystals to intracellular compartments, predominantly LDs (Movie 2). This observation was confirmed by quantitative analysis, which showed that the LD volume fraction per cell increased from 0.0046 ± 0.0003 to 0.0113 ± 0.0005 following M*β*CD treatment (Figure 5C). In addition, LDs in M*β*CD-treated cells contained significantly higher TF-Chol fluorescence (149.3 ± 7.5) compared to untreated cells (67.3 ± 8.0), indicating that cholesterol released from dissolving crystals is efficiently stored in LDs (Figure 5D).

**Figure 5:**
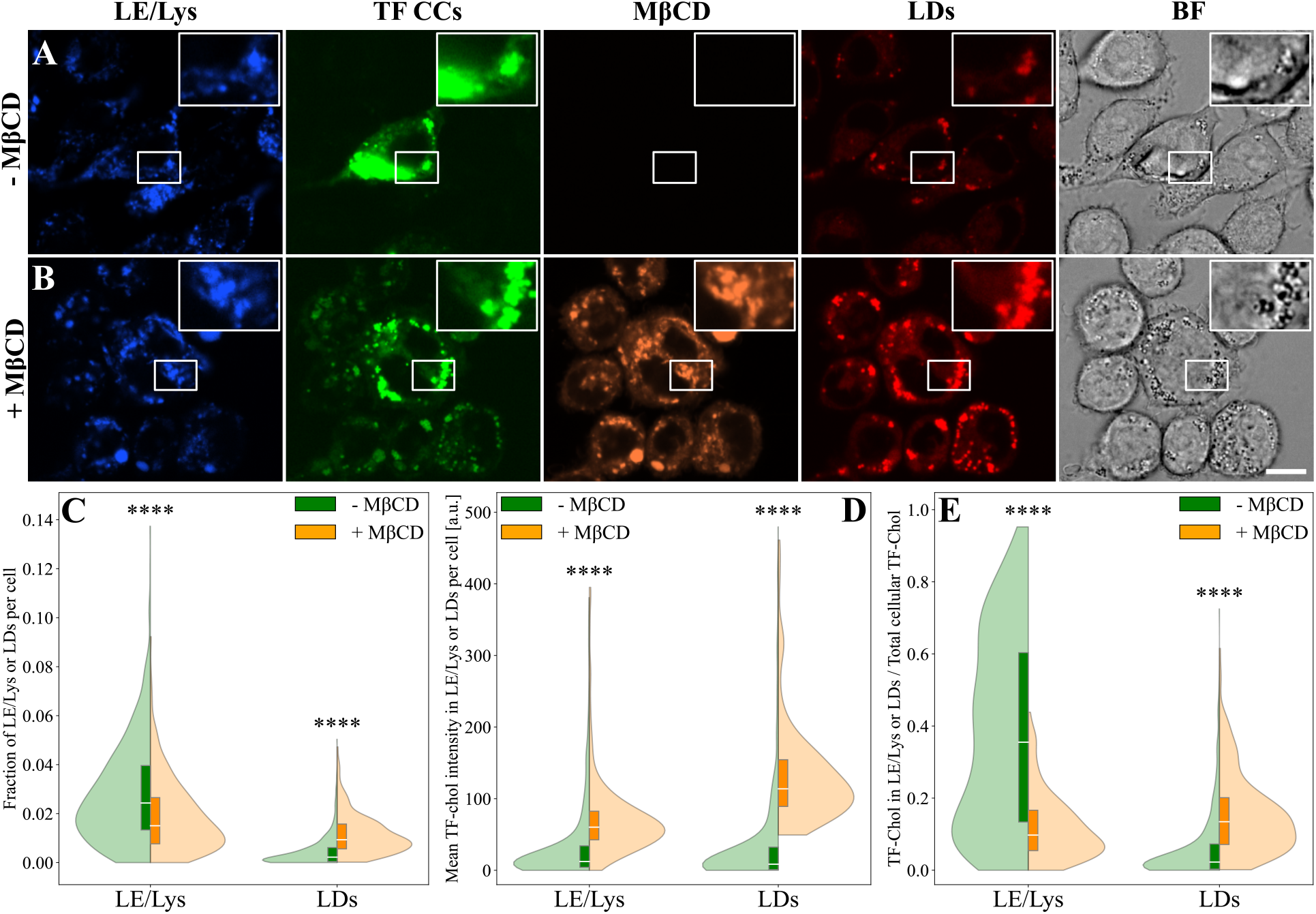
Cyclodextrin promotes dissolution of TF CCs in macrophages. Macrophages were incubated with TF CCs for 5 h to allow crystal uptake, washed, and either maintained in fresh medium (A) or treated overnight with a mixture of fluorescent and non-fluorescent M*β*CD (1:10, final concentration of 1 mM) (B). Scale bar, 10 µm. Quantification was performed from five independent experiments for cells treated with TF CCs (n = 488 cells) and three independent experiments for cells with TF CCs + M*β*CD (n = 265 cells). LE/Lys and LD volume fractions, calculated as the number of organelle-positive voxels relative to the total number of voxels within the cell mask, in untreated cells (green) or treated cells (orange) (C). Mean TF-Chol fluorescence intensity within LE/Lys or LDs per cell (D). Fraction of total cellular TF-Chol fluorescence intensity detected within LE/Lys or LDs (E). Statistical significance was determined using Mann-Whitney U test: p = 2.50 *×* 10*^−^*^10^ (C, LE/Lys), 3.68 *×* 10*^−^*^41^ (C, LDs), 2.78 *×* 10*^−^*^46^ (D, LE/Lys), 1.31 *×* 10*^−^*^64^ (D, LDs), 2.02 *×* 10*^−^*^39^ (E, LE/Lys), and 9.47 *×* 10*^−^*^43^ (E, LDs).

Fluorescent cyclodextrin was internalized by macrophages and localized to LE/Lys, suggesting that cyclodextrin can access intracellular LE/Lys and may thereby facilitate the dissolution of intracellular CCs, in addition to its proposed effect at the plasma membrane.^18^ Additionally, we find that the fluorescent cyclodextrin shows no accumulation at intracellular TF CCs (Figure S5). Notably, cyclodextrin treatment also altered the LE/Lys volume fraction, which decreased from 0.028 ± 0.0009 to 0.019 ± 0.0009 following M*β*CD treatment (Figure 5C). Despite this reduction, the mean TF-Chol fluorescence intensity within the LE/Lys increased from 41.3 ± 4.4 to 74.4 ± 4.1, consistent with transient accumulation of crystal-derived cholesterol within LE/Lys following crystal dissolution (Figure 5D). However, the fraction of total cellular TF-Chol fluorescence detected within LE/Lys decreased markedly (0.38 ± 0.012 versus 0.12 ± 0.0057), whereas the fraction detected within LDs increased from 0.0625 ± 0.0044 to 0.15 ± 0.0067 (Figure 5E). Together, these findings indicate that M*β*CD promotes the dissolution of intracellular CCs and redistributes the released cholesterol from the lysosomal compartment toward storage in LDs. This redistribution may represent a protective mechanism that reduces the burden of crystalline cholesterol while facilitating intracellular cholesterol sequestration.

### Uptake and Dissolution of Dehydroergosterol Crystals by Macrophages

While being a useful tool to label CCs and study their uptake and processing by macrophages, TF-Chol’s physicochemical and biological properties differ significantly from those of cholesterol due to the attached BODIPY moiety.^24,36^ To rule out that the observed effects of M*β*CD were specific to TF-Chol-labelled crystals, we generated crystals of the intrinsically fluorescent sterol analog DHE and compared the emission spectra before and after treatment with M*β*CD. Upon excitation at 328 nm, monomeric DHE in ethanol exhibited only one emission peak at 375 nm, whereas crystalline DHE displayed two additional peaks around 400 and 425 nm (Figure S6A,B), consistent with previous observations.^21^

Monitoring DHE crystals before and after addition of M*β*CD (1 mM) revealed a rapid shift in the emission profile toward the monomeric DHE profile. Specialized UV-sensitive widefield fluorescence microscopy further revealed that fluorescent cyclodextrin associated with DHE crystals, supporting a direct interaction between cyclodextrin and the crystal surface *in vitro* (Figure S7). After 30 min, the majority of the fluorescence was detected at 375 nm, and the crystal-associated peaks at 400 and 425 nm were markedly reduced (Figure 6B). These findings indicate that crystalline DHE is self-quenched and that crystal dissolution by M*β*CD restores the fluorescence characteristics of monomeric DHE.

**Figure 6:**
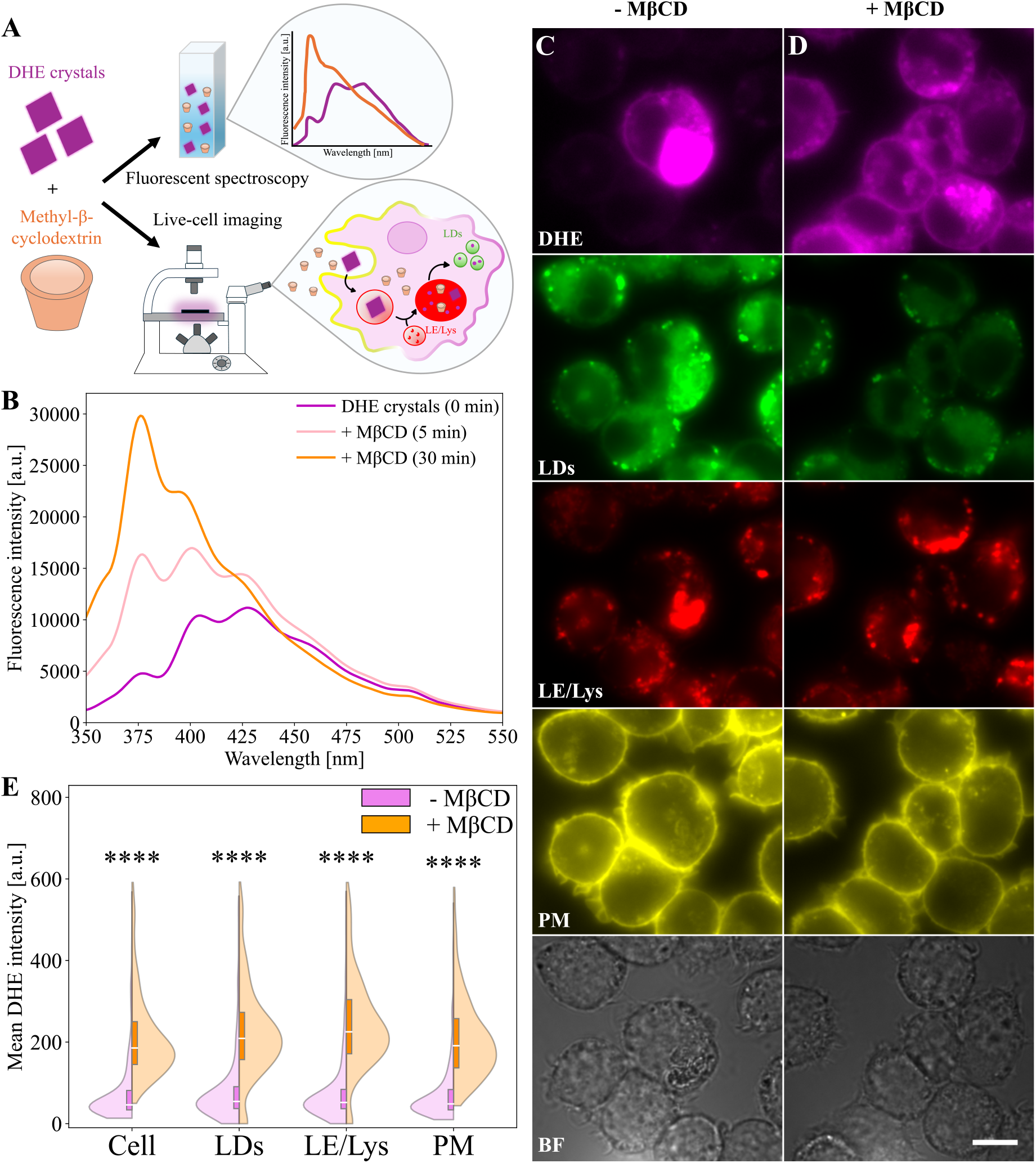
Cyclodextrin promotes dissolution of DHE crystals. Schematic overview of the experimental design (A). Emission spectra of DHE crystals (0.8 mg) following the addition of M*β*CD (1 mM), shown as the mean of two independent experiments (B). Macrophages were incubated with DHE crystals (0.08 mg) for 5 h, washed, and either maintained in fresh LPDS medium (C) or treated overnight with M*β*CD (1 mM) (D). Scale bar, 10 µm. Quantification was performed from three independent experiments (553 cells in the DHE crystal group and 518 cells in the M*β*CD-treated group). Mean DHE fluorescence intensity in the whole cell (p = 3.14*×*10*^−^*^93^), lipid droplets (LDs, p = 4.17*×*10*^−^*^81^), late endosomes/lysosomes (LE/Lys, p = 1.34 *×* 10*^−^*^118^), and the plasma membrane (PM, p = 1.82 *×* 10*^−^*^115^) for cells containing only DHE crystals (violet) or following M*β*CD treatment (orange) (E).

Having established that M*β*CD dissolves DHE crystals *in vitro*, we next investigated whether the same process occurs in cells. Macrophages were incubated with DHE crystals for 5 h to allow crystal uptake before treating cells overnight with M*β*CD. Cells were subsequently imaged using two specialized UV-sensitive widefield fluorescence microscope set ups, enabling both 2D (Figure 6C,D), and 3D imaging, respectively (Figure S8). Using fluorescent markers for the LDs, LE/Lys, and the PM, we quantified the distribution of DHE in these cellular compartments (Figure 6E). M*β*CD treatment significantly increased DHE fluorescence throughout the cell from 98.8 ± 7.6 to 225.3 ± 6.2, including within LDs (127.1 ± 11.5 to 248.6 ± 8.0), LE/Lys (162.2 ± 17.7 to 284.4 ± 9.7), and the PM (79.9 ± 4.4 to 212.7 ± 4.8) (Figure 6E). Consistent with the fluorescence emission spectra, the increase in intracellular DHE fluorescence following M*β*CD treatment is compatible with dequenching of DHE as the crystals dissolve. In contrast to the TF CCs, however, M*β*CD treatment did not significantly alter the fraction of cell area occupied by LDs or LE/Lys (Figure S6C). A possible explanation for this difference is that CCs doped with TF-Chol mostly contain cholesterol (i.e. in a 8:1 ratio, see Materials and Methods), while DHE crystals are made from the intrinsically fluorescent sterol, only. Cholesterol is a much better substrate for ACAT than DHE,^37^ indicating that most cholesterol but not DHE gets esterified after crystal dissolution, explaining the higher fraction of LDs in cells treated with TF-CCs compared to macrophages treated with DHE crystals. The distribution of DHE between LDs and LE/Lys was similar to that observed for TF-Chol, with a reduced fraction of total cellular DHE fluorescence detected within LE/Lys and an increased fraction detected within LDs following M*β*CD treatment (Figure S6D). Together, these findings show that M*β*CD-mediated crystal dissolution is not unique to TF-Chol-labeled crystals but is also observed using the minimally modified sterol analog DHE, which supports that cyclodextrin promotes dissolution of cholesterol crystals rather than redistribution of the fluorescent probe.

## Discussion

Formation of CCs is a hallmark of atherogenesis, but the mechanisms by which these crystals can be cleared are poorly understood.^5^ In this study, we show that macrophages efficiently internalize large and medium-sized CCs, while they also employ exophagy for degradation of large CCs. Importantly, we find that intact lysosomal function is key for both the intra- and extracellular digestion of CCs. Using novel 3D correlative fluorescence and soft X-ray microscopy combined with 3D imaging assays of crystal processing, we demonstrate that macrophages internalize CCs of varying sizes and shapes (e.g., needles and plates) and dynamically engage with the crystal surface. These results extend earlier findings employing electron and light microscopy as well as SXT to show that CCs of needle- and plate-like shapes can form in cholesterol-loaded macrophages.^6,11,12,38–40^ Similarly, needle- and plate-shaped CCs have been observed in the extracellular space of atherosclerotic lesions using transmission electron microscopy, polarized light microscopy, and cryo-focused ion beam milling electron microscopy (cryo-FIB-SEM).^7,10,41^ We find that engagement of macrophages with large CCs recruits LE/Lys to the crystal surface, where lysosomal enzymes can lead to degradation of the CCs. This process continues inside cells and requires intact acidification via the vacuolar proton pump, as does the subsequent targeting of liberated TF-Chol to LDs. Upon inhibition of NPC1 function using U18666A, export of cholesterol liberated from CCs is reduced, demonstrating that lysosomal sterol export depends on the canonical NPC1/NPC2 pathway. The crystal-derived cholesterol exported from LE/Lys is targeted to LDs, as we show by 3D live-cell imaging of TF-Chol, thereby supporting indirect findings using cryo-FIB-SEM by Addadi and co-workers.^40^ The expansion of LDs following U18666A treatment could be due to reduced autophagy of LDs (lipophagy) in cells with inhibited NPC1 activity. Indeed, increased abundance of LDs has also been found in NPC1-deficient cells, as observed in Niemann-Pick type C disease.^42^ Furthermore, lipophagy has been shown to be a major clearance pathway of excess cholesterol in macrophage foam cells.^43^ Lysosomal exocytosis and extracellular acidification have previously been shown to be necessary for exophagy of agLDL by macrophages,^3^ and the acidic environment of atherosclerotic plaques may therefore play a key role in the clearance capacity of not only agLDL but also of CCs by macrophages and other cells.^44,45^

It has previously been shown that CDs can dissolve cholesterol deposits and thereby promote regression of atherosclerosis.^16^ CDs can also reduce inflammatory responses of macrophages induced by CCs and reduce the overall uptake of CCs by these cells.^46,47^ On the subcellular level, CDs increase the formation of cholesteryl esters and oxysterols from CCs in macrophages.^16^ Our results show that CDs can be internalized and promote CC dissolution from within LE/Lys, thereby increasing the formation of LDs and accelerating crystal degradation. That CDs act from inside endo-lysosomes has been demonstrated in another context, namely the restoration of normal cholesterol levels in lysosomal storage diseases, such as Niemann-Pick type C disease.^18^ Using the unique spectral changes of the intrinsically fluorescent sterol DHE upon incubation with M*β*CD, we were able to directly monitor the solubilization of the sterol crystals by CDs. Such *in vitro* experiments were complemented by the observation of efficient DHE trafficking from ingested crystals inside LE/Lys to the PM and LDs, further substantiating the main role of lysosomal processing of CCs and its enhancement by CDs in macrophages.

## Materials and Methods

### Materials

The murine macrophage cell line J774A.1 (J774,#TIB-67) was purchased from the American Type Culture Collection (ATCC). T25 Nunc™ EasyFlask™ culture flasks (#156367), Heat-inactivated Fetal Bovine Serum (HI FBS, #A5209502), Dextran Cascade Blue, 10,000 MW (CB, #D1976), Dextran, Tetramethylrhodamine, 70,000 MW (Rh-dex, #D1819), HCS LipidTOX Deep Red (#H34477), Dextran tetramethylrhodamine and biotin, 10,000 MW, mini-ruby (#D3312), NeutrAvidin protein (#31000), Hoechst 33342 (#62249), CellMask Deep Red (#C10046), and BODIPY 493/503 (#D3922) were purchased from Thermo Fisher Scientific. Dulbecco’s Modified Eagle Medium (DMEM, #D6429), Penicillin-streptomycin solution (P/S, #P4333), Lipoprotein Deficient Serum from human plasma (LPDS, #S5519), U18666A (#U3633), Bafilomycin A1 (Baf-A1, #B1793), Methyl-*β*-cyclodextrin (M*β*CD, #332615), biotin (#B4501), Ergosta-5,7,9(11),22-tetraen-3*β*-ol (DHE, #E2634), and Sodium oleate (#O7501) were purchased from Sigma-Aldrich. Glass-bottom 35 mm microscope dishes (#TKO-P351184-408) were purchased from MatTek, and µ-Slide 8 Well high chambered coverslips (#80807) were purchased from Ibidi. Cholesterol (#197144) and TopFluor-Cholesterol (TF-Chol, #810255P) were purchased from Avanti Polar Lipids. Rhodaminyl-thioureido-RAME*β* (#CY-R-2004.1) was obtained from CycloLab. Grids, R 2/2 AU 200 mesh (#Q92210) were purchased from Quantifoil. Cholesterol-PEG-biotin, MW 1,000 (#BP-25772) was purchased from Broadpharm.

### Cell culture

J774 macrophages were cultured in T25 flasks and grown in DMEM supplemented with 10 % HI FBS and 1 % P/S solution at 37 °C, 5 % CO_2_, and 95 % humidity. Cells were passaged every 3 days by scraping and split in a ratio of 1:5. For microscopy experiments, cells were plated in 35 mm glass-bottom microscope dishes or 8 Well high chambered coverslips and cultured until approximately 80 % confluency before imaging.

### Formation of cholesterol crystals

CCs containing TF-Chol (TF-CCs) were prepared by transferring 0.08 mg cholesterol from a chloroform stock solution to a glass vial, followed by evaporation of the solvent under a stream of nitrogen. The dried cholesterol was redissolved in absolute ethanol (abs. EtOH), and 0.01 mg TF-Chol from an abs. EtOH stock solution was added to a final volume of 1 mL. Sterile water (10 % of the total volume) was added, and the solution was mixed before incubation at room temperature in the dark for 15 min to allow crystal formation. The solvent was subsequently evaporated at 40 °C under nitrogen. The resulting crystals were crushed using a spatula, resuspended in 1 mL DMEM supplemented with 10 % LPDS and 68.8 nM sodium oleate (DMEM/LPDS), and added directly to cells.

For the lysosomal exocytosis assay, CCs were prepared with the addition of cholesterol-PEG-biotin and NeutrAvidin. In short, 0.08 mg cholesterol, 0.005 mg TF-Chol, and 0.005 mg cholesterol-PEG-biotin (1 % w/w) were transferred to a glass vial. Sterile water was added to 10 % of the final volume, as described above. NeutrAvidin was added in excess to the crystals/medium solution to ensure binding to all cholesterol-PEG-biotin molecules while leaving unoccupied biotin-binding sites available for subsequent binding of biotinylated dextran.

DHE crystals were prepared using either 0.08 mg DHE for cell-based experiments or 0.8 mg DHE for fluorescence spectroscopy experiments. DHE was dissolved in abs. EtOH (1.97 mg/mL stock solution), and crystals were generated following the same procedure described for TF-Chol crystals.

### Cholesterol crystal uptake in macrophages

Cells were washed twice with PBS before incubation with TF CCs in DMEM/LPDS medium for 5 h. Following crystal uptake, cells were washed twice with PBS to remove non-cell-associated CCs. For crystal dissolution experiments, cells were incubated overnight with a mixture of fluorescent and non-fluorescent M*β*CD (1:10) (1 mM) in DMEM/LPDS. To inhibit NPC1-mediated cholesterol export from LE/Lys, cells were treated overnight with U18666A (5 µM). To inhibit lysosomal acidification, cells were pretreated with Baf-A1 (50 nM) for 1 h before TF CCs exposure.

For visualization of late endosomes/lysosomes (LE/Lys), cells were incubated overnight with either Rh-dextran (25 µg/mL) or Cascade Blue dextran (125 µg/mL). Before imaging, cells were washed twice with Medium 1 (M1) (containing 150 mM NaCl, 5 mM KCl, 1 mM CaCl_2_, 1 mM MgCl_2_, 5 mM glucose and 20 mM HEPES, pH 7.4). Lipid droplets (LDs) were labelled with HCS LipidTOX Deep Red (0.5X) or BODIPY (20 ng/mL) for 5 min before imaging. For staining the plasma membrane, cells were incubated with CellMask Deep Red (1 µg/mL) for 5 min before imaging.

### Lysosomal exocytosis assay with cholesterol crystals

The lysosomal exocytosis assay is a well-established method for studying macrophage digestion of agLDL and apoptotic cells.^26–28,34^ To adapt this assay for CCs, J774 macrophages were plated in 8-well high-chambered coverslips (approximately 90,000 cells/well) and grown overnight in complete medium. Biotin-tetramethylrhodamine-dextran (1 mg/mL) was added overnight for delivery to the lysosomes. The following day, cells were washed three times with complete medium, and dextran was chased for 3 h to ensure delivery to the lysosomes. Cells were afterwards washed three times, and for drug inhibition experiments, incubated for 1 h in DMEM/LPDS medium before adding NeutrAvidin-biotin-TF CCs for 90 min. During this time, any biotin-dextran released by lysosomal exocytosis toward the NeutrAvidin-biotin-TF-CCs was captured at the crystal surface through biotin-NeutrAvidin binding. Cells were subsequently incubated for 15 min with excess biotin (200 µM in DMEM without supplements) to block any remaining unoccupied biotin-binding sites on NeutrAvidin. Finally, cells were fixed for 15 min at room temperature (RT) in 0.5 % paraformaldehyde (PFA) in PBS containing Hoechst 33342 (5 µM) and afterwards stained with HCS LipidTOX Deep Red (1:1000).

### Confocal fluorescence microscopy

A Nikon A1 confocal microscope with a Nikon Ti-1 LFOV body and a 60X (Plan, Apo, VC, Water, NA 1.2, WD 0.31 mm) objective was used for imaging. Images were acquired using the Galvano scanner with two-frame averaging and a Z-step size of 0.5 µm (45 optical sections per stack). CB was imaged using a 405 nm laser (0.40 % laser power, gain 80, offset 12), TF-Chol was imaged using a 488 nm laser (0.20 % laser power, gain 1, offset 5), RAME*β* and Rh-dextran were imaged using a 561 nm laser (0.5 % laser power (for RAME*β*) and 2 % (for Rh-dextran), gain 5, offset 10), and LipidTOX was imaged using a 637 nm laser (0.5 % laser power, gain 70, offset 10). For time series imaging, an Okolab microscope stage incubator was used, allowing the cells to be imaged at 37 °C and 5 % CO_2_. For the lysosomal exocytosis assay, images were acquired as 2×2 or 3×3 tilted scans with 15 % overlap between adjacent tiles. For each tiled scan, a Z-stack spanning a total depth of 10 µm was acquired with a 1 µm step size.

### UV fluorescence microscopy

UV-sensitive epifluorescence microscopy was carried out on two different microscope systems. The first system consisted of a Leica DMIRBE microscope with a 63 × 1.4 NA oil immersion objective (Leica Lasertechnik GmbH) with a Lambda SC smart shutter (Sutter Instrument Company) as illumination control. Images were acquired with an Andor Ixon blue EMCCD camera operated at -75 °C and controlled using Solis software. The microscope contained a 10× extra magnification lens in the emission light path, resulting in a final pixel size of 193 nm for the 63× objective. DHE was imaged using a UV-adapted filter cube obtained from Chroma Technology (Corp., Brattleboro, VA, USA) with a 335 nm (20 nm bandpass) excitation filter, 365 nm dichromatic mirror, and 405 nm (40 nm bandpass) emission filter. A bleach stack with 100 frames was recorded with an exposure time of 400 ms. BODIPY was imaged using a fluorescein filter cube with 480 nm (40 nm bandpass) excitation filter, 505 nm longpass dichromatic mirror, and 527 nm (30 nm bandpass) emission filter. Rh-dextran was imaged using a Rhod Et filter cube with a 530 nm (30 nm bandpass) excitation filter, a 580 nm dichromatic mirror, and a 590 nm emission filter. Images were taken with an exposure time of 0.1 s. The CellMask dye was imaged using the IR channel. The infrared filter cube used an excitation filter of 620 nm (20 nm bandpass), a 660 nm dichromatic mirror, and a 700 nm (75 nm bandpass) emission filter. Images were taken with an exposure time of 0.4 s. Additionally, UV-sensitive epifluorescence imaging was performed using a Leica Microsystems Leica DMi8 inverted microscope equipped with a Leica K8 camera and a 63×/1.40–0.60 oil immersion objective lens (#11506349). Illumination was provided by a pE-400max LED light source (#11504265, Leica Microsystems) and a 7UV illumination source (UVICO-2-DUV), consisting of a fibre optically coupled UVC–VIS light source with a 200 W lamp, a power supply (110–240 V, 50/60 Hz), bandwidth selections of 220–400 nm and 400–700 nm, an LLG-3/2 DUV liquid light guide (3 mm diameter, 2 m length). DHE fluorescence was imaged using a RappOpto special filter cube with excitation at 330/20 nm, a dichroic long-pass filter at 360 nm (DCLP 360), and emission at 405/40 nm. Rh-dextran fluorescence was imaged using a Filter Cube LED 530 (#11525342, size P) with a 525/50 nm excitation, a dichroic filter at 560 nm, and emission at 605/70 nm. CellMask fluorescence was imaged using a Filter Cube Y5 (#11525312, size P) with excitation at 620/60 nm, a dichroic filter at 660 nm, and emission at 700/75 nm.

### Fluorescence spectroscopy of DHE crystals

The emission spectra of DHE crystals were obtained using an ISS Chronos BH spectrofluorometer (Urbana-Champaign, IL). The excitation wavelength was set to 328 nm, and emission was recorded from 350 to 550 nm. For all measurements, a slit width of 0.5 mm (excitation and emission path) with no polarization was used. For fluorescence spectroscopy, DHE crystals were prepared using 0.8 mg DHE, corresponding to a 10-fold higher concentration than that used for cell-based experiments, to increase the fluorescence signal. The crystals were subsequently resuspended in M1 medium, unless otherwise stated. Emission spectra were recorded immediately before the addition of M*β*CD (final concentration of 1 mM) and after 5 and 30 min of incubation. Each spectrum represents the mean from two independent measurements unless otherwise stated. Spectral data were processed in Python. Individual spectra were smoothed using a Savitzky–Golay filter before averaging.

### Image analysis of fluorescence microscopy data

#### Cell segmentation using Cellpose

Cell segmentation was performed using the deep learning-based Cellpose algorithm, as described in.^48^ For 3D confocal image stacks, cells were segmented based on the LipidTOX fluorescence channel. Before segmentation, images were blurred by applying a 3D Gaussian blur (*σ* = 6) to reduce image noise. Segmentation was performed using the built-in Cytoplasm 2.0 model (cyto2) with an estimated cell diameter of 120 pixels, a minimum object size of 300 pixels, an anisotropy factor of 2.0, a cellprob threshold of -6, and a flow threshold of 0.4 with 3D segmentation enabled. Image preprocessing and segmentation were fully automated and performed in batch mode using in-house Python scripts.

For 2D images acquired by the UV-sensitive epifluorescence microscope, cells were segmented based on the CellMask fluorescence images. Segmentation was performed using the Cytoplasm 2.0 model (’cyto2’) with an estimated cell diameter of 100 pixels and a flow threshold of 0.4.

#### Quantitative analysis of confocal images

For 3D confocal image analysis, late endosomes/lysosomes (LE/Lys) were segmented in each optical section using intensity-based thresholding following background correction. Uneven background fluorescence was removed independently from each Z-slice by morphological opening using a disk-shaped structuring element with a radius of 10 pixels. The resulting background-corrected 3D image stack was subsequently smoothed using a Gaussian filter (*σ* = 1). LE/Lys were segmented using a slice-specific intensity threshold corresponding to seven times the mean intensity of the background-corrected Z-slice.

TF-CCs were segmented by intensity-based thresholding following 3D morphological background correction using a spherical structuring element with a radius of 10 pixels and Gaussian smoothing (*σ* = 1). A single threshold factor was applied to all images within each experimental condition, while threshold factors were adjusted between conditions to account for differences in fluorescence intensity and crystal morphology. The resulting CC masks were used to exclude crystal-associated TF-Chol fluorescence from subsequent intracellular analyses.

LDs were manually annotated slice-wise in five near-fully labeled volumes, and these annotations were used to train a hybrid 3D-to-2D U-Net.^49^ To compensate for the limited amount of annotated data, training pairs were augmented on the fly with random flips along both in-plane axes (*p* = 0.5 each), random multiples of 90*^◦^*in-plane rotation, and random elastic deformation (*p* = 0.9). Image intensities were further perturbed by random brightness and contrast jitter (*±*10%) and additive positive-only Gaussian noise with a standard deviation of up to 2% of the image standard deviation. The network was optimized with the sum of a soft Dice loss and a focal loss (*γ* = 2, *α* = 0.25). Unannotated voxels, including those introduced at the image borders by elastic deformation, were assigned an ignore label and excluded from the loss, allowing the network to be trained on incompletely annotated volumes. Given the relatively simple and stereotyped appearance of LDs, this scheme was sufficient to train an accurate model from five annotated volumes, corresponding to a soft Dice coefficient greater than 0.95. Segmentations were additionally inspected visually by an expert annotator and judged to be of at least human-level quality. The trained model was subsequently applied to the remaining images to generate LD masks for quantitative analysis.

Quantitative analysis was performed on a per-cell basis across the confocal Z-stacks. For each segmented cell, cell volume and the volume fraction occupied by LE/Lys and LDs were determined from the corresponding binary organelle masks. The organelle volume fraction was calculated as the number of organelle-positive voxels within a cell divided by the total number of voxels within the corresponding 3D cell mask. TF-Chol fluorescence was quantified after exclusion of pixels with fluorescence intensities below 60 to reduce background signal. For each cell, the total TF-Chol fluorescence within the LE/Lys or LD mask and the total TF-Chol fluorescence within the cell mask were determined across the Z-stack. Mean TF-Chol fluorescence intensity within each compartment was calculated as the summed TF-Chol fluorescence within the corresponding organelle mask divided by the number of organelle-positive voxels. The fraction of total cellular TF-Chol fluorescence detected within LE/Lys or LDs was calculated as the summed TF-Chol fluorescence within the respective organelle mask divided by the total TF-Chol fluorescence within the corresponding cell mask. Only cells with a segmented volume greater than 100,000 voxels were included in the quantitative analyses.

#### Lysosomal exocytosis analysis

Image analysis of the lysosomal exocytosis assay was performed using an automated 2D analysis workflow implemented in Python. Cell segmentation was performed from maximum-intensity projections of confocal Z-stacks using Cellpose as earlier described. Cells were segmented in 2D using the Cytoplasm 2.0 model (cyto2) with an estimated cell diameter of 150 pixels and a flow threshold of 0.4. CCs were segmented from the crystal fluorescence channel using intensity-based thresholding. A maximum-intensity projection of the resulting binary 3D crystal mask was generated for subsequent 2D analysis. For quantification of Rh-dextran fluorescence, a constant background value of 500 intensity units was subtracted from the Rh-dextran channel, and negative values were set to zero. A maximum-intensity projection was subsequently generated from the background-corrected Z-stack. Rh-dextran association with cholesterol crystals was quantified as the number of pixels within the projected crystal mask with Rh-dextran fluorescence above the applied background threshold. To quantify cellular Rh-dextran fluorescence, the mean Rh-dextran fluorescence intensity was calculated separately within each Cellpose-derived cell mask, and the resulting per-cell values were averaged for each image.

#### Quantitative analysis of widefield images

For 2D images acquired by the UV-sensitive epifluorescence microscope, the plasma membrane (PM) regions were generated from the Cellpose-derived cell masks. For each segmented cell, the cell mask was morphologically eroded using a disk-shaped structuring element with a radius of 2 pixels. The eroded mask was subtracted from the original cell mask to generate a 2-pixel-wide region along the inner cell boundary, which was defined as the PM region. The resulting PM masks were subsequently used for quantification of DHE fluorescence intensity in the PM. The LE/Lys and LDs were segmented from the Rh-dextran and BODIPY fluorescence channels, respectively. For both compartments, uneven background fluorescence was estimated by morphological opening with a disk-shaped structuring element and subtracted from the original image. The background-corrected images were subsequently smoothed using a Gaussian filter (*σ* = 1). LE/Lys were segmented using a disk radius of 5 pixels and an intensity threshold corresponding to six times the mean intensity of the background-corrected image, whereas the LDs were segmented using a disk radius of 4 pixels and a threshold corresponding to eight times the mean intensity of the background-corrected image. Pixels exceeding the respective thresholds were classified as belonging to the corresponding compartment, and the resulting binary masks were used for subsequent fluorescence intensity and compartment-based analyses.

For each segmented cell, the area fraction occupied by each compartment was calculated as the number of organelle-positive pixels within the cell divided by the total number of pixels within the corresponding cell mask. Mean DHE fluorescence intensity within each compartment was calculated as the summed DHE fluorescence within the corresponding compartment mask divided by the number of compartment-positive pixels. The fraction of total cellular DHE fluorescence detected within each compartment was calculated as the summed DHE fluorescence within the corresponding compartment mask divided by the total DHE fluorescence within the cell mask.

All microscopy experiments included at least three independent experiments. Statistical analyses were performed in Python (Jupyter Notebook, version 7.3.2). Data are presented as mean ± SEM unless otherwise stated. Comparisons between independent groups were performed using the two-sided Mann–Whitney U test. The number of independent experiments (N) and cells analyzed (n) are indicated in the corresponding figure legends. A *p*-value *<* 0.05 was considered statistically significant.

#### Cell preparation for Soft X-ray microscopy

Quantifoil grids were autoclaved before use, coated with Poly-D-Lysine for 1.5 h at 37 °C, and washed twice with sterile water before seeding J774 macrophages onto the grids. Cells were cultured on the grids for two days before adding TF CCs in 1 mL DMEM/LPDS overnight. Cells were washed three times with PBS before fixation with 2 % PFA, pH 7.36, for 15 min on ice. The cells were subsequently cryopreserved by vitrification using plunge freezing. TEM grids bearing the cells were rapidly plunged into liquid ethane cooled by liquid nitrogen using a Leica EM GP2 plunge freezer. The blotting time was set to 6 s, the relative chamber humidity to 80 %, and the liquid ethane temperature to -182 °C.

#### Soft X-ray microscopy

Soft X-ray tomography (SXT) imaging was performed using two different microscope setups. Synchrotron-based SXT images were acquired at the U41 TXM beamline at BESSY II, Helmholtz Zentrum Berlin (HZB), using 25 nm zone plates over a tilt range of 120–125° with 1° tilt steps at 510 eV. The exposure time per projection was 6–12 s, with a pixel size of 9.8 nm. Tomographic projections were aligned using Bsoft and reconstructed by filtered back-projection using Tomo3D.^50,51^

Alternatively, SXT images were acquired using the laboratory-based SXT-100 microscope (SiriusXT, Dublin, Ireland).^31^ SXT tilt series were acquired over a total angular range of 121*^◦^* (−60*^◦^* to + 60*^◦^*) with 1*^◦^* tilt increments at an X-ray photon energy of 453 eV. The exposure time per projection ranged from 60 to 180 s, depending on sample thickness and tilt angle, with longer exposures used for thicker samples and higher tilt angles. The tilt series were aligned using a custom-written tilt-series alignment program that employed a combination of path tracking and fiducial-marker tracking, with scale-free feature detection used to detect and initialise virtual markers, as previously described.^52^ The aligned tilt series were subsequently reconstructed using the simultaneous iterative reconstruction technique (SIRT) implemented in Tomo3D, with 50 iterations.^51^

## Acknowledgments

We thank Frederick R. Maxfield (Weill Medical College of Cornell University, New York, USA) for his support with the lysosomal exocytosis assay and helpful discussions. This work was supported by the Danish Council for Independent Research (grant agreement No. 2032-00136B) to Daniel Wüstner. Ultraviolet (UV) 3D imaging was performed using a Leica microscope, funded by the Carlsberg Foundation (grant agreement No. CF24-1904). Confocal imaging was performed at the Danish Molecular Biomedical Imaging Center (DaMBIC, University of Southern Denmark), supported by the Novo Nordisk Foundation (NNF) (grant agreement No. NNF18SA0032928). Development of the tilt-series alignment program was funded by the European Union’s Horizon 2020 research and innovation programme through the CoCID project (grant agreement No. 101017116). We thank the Helmholtz-Zentrum Berlin (HZB) für Materialien und Energie for the allocation of synchrotron radiation beamtime.

## Author Competing Interests

The authors declare that they have no known competing financial interests or personal relationships that could have appeared to influence the work reported in this paper.

## Supporting Information

**Figure S1:**
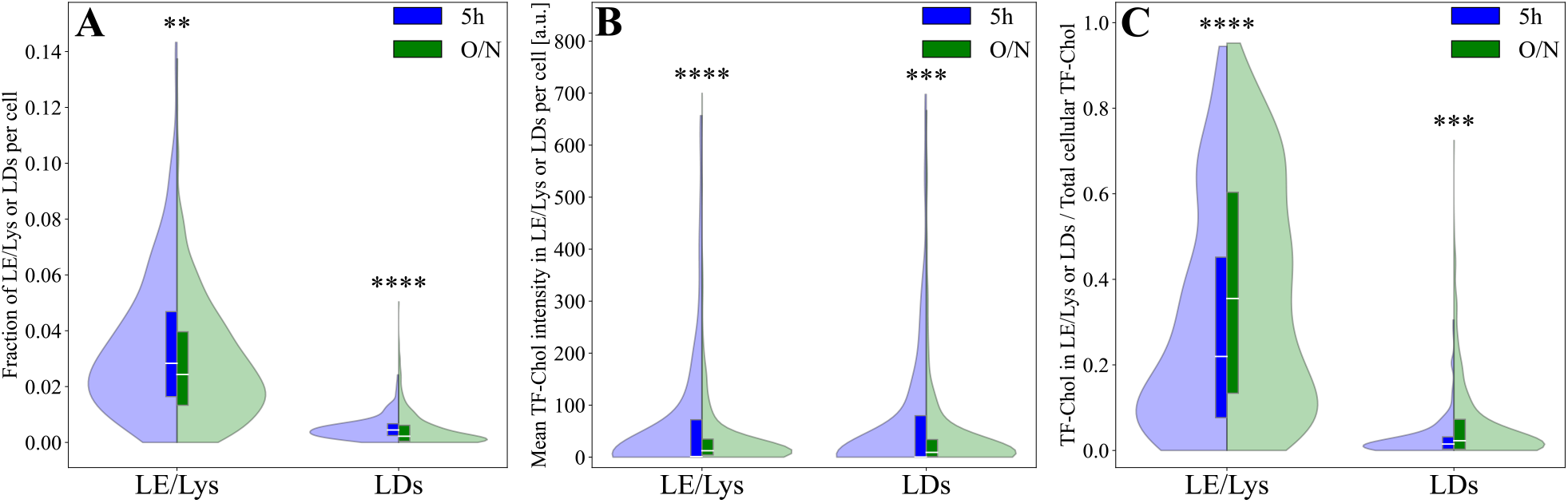
Intracellular distribution of TF-Chol following uptake of TF CCs in J774 macrophages after 5 h or overnight (O/N) incubation. The LE/Lys and LD volume fraction were quantified after 5 h (blue) or overnight (O/N, green) incubation with TF CCs (A). Mean TF-Chol fluorescence intensity within LE/Lys and LDs per cell was quantified after 5 h and O/N incubation (B). The fraction of total cellular TF-Chol fluorescence detected within LE/Lys or LDs was quantified as the TF-Chol fluorescence within the respective organelle mask divided by the total TF-Chol fluorescence within the cell (C). Quantification was performed from three independent experiments for 5 h TF CCs-treated cells (n = 219 cells), and five independent experiments for cells with TF CCs O/N treatment (n = 488 cells). Statistical significance was assessed using Mann-Whitney U test (p = 1.58 *×* 10*^−^*^3^ (A, LE/Lys), 2.61 *×* 10*^−^*^8^ (A, LDs), 3.82 *×* 10*^−^*^8^ (B, LE/Lys), 1.38 *×* 10*^−^*^4^ (B, LDs), 1.81 *×* 10*^−^*^5^ (C, LE/Lys) and 2.74 *×* 10*^−^*^4^ (C, LDs).

**Figure S2:**
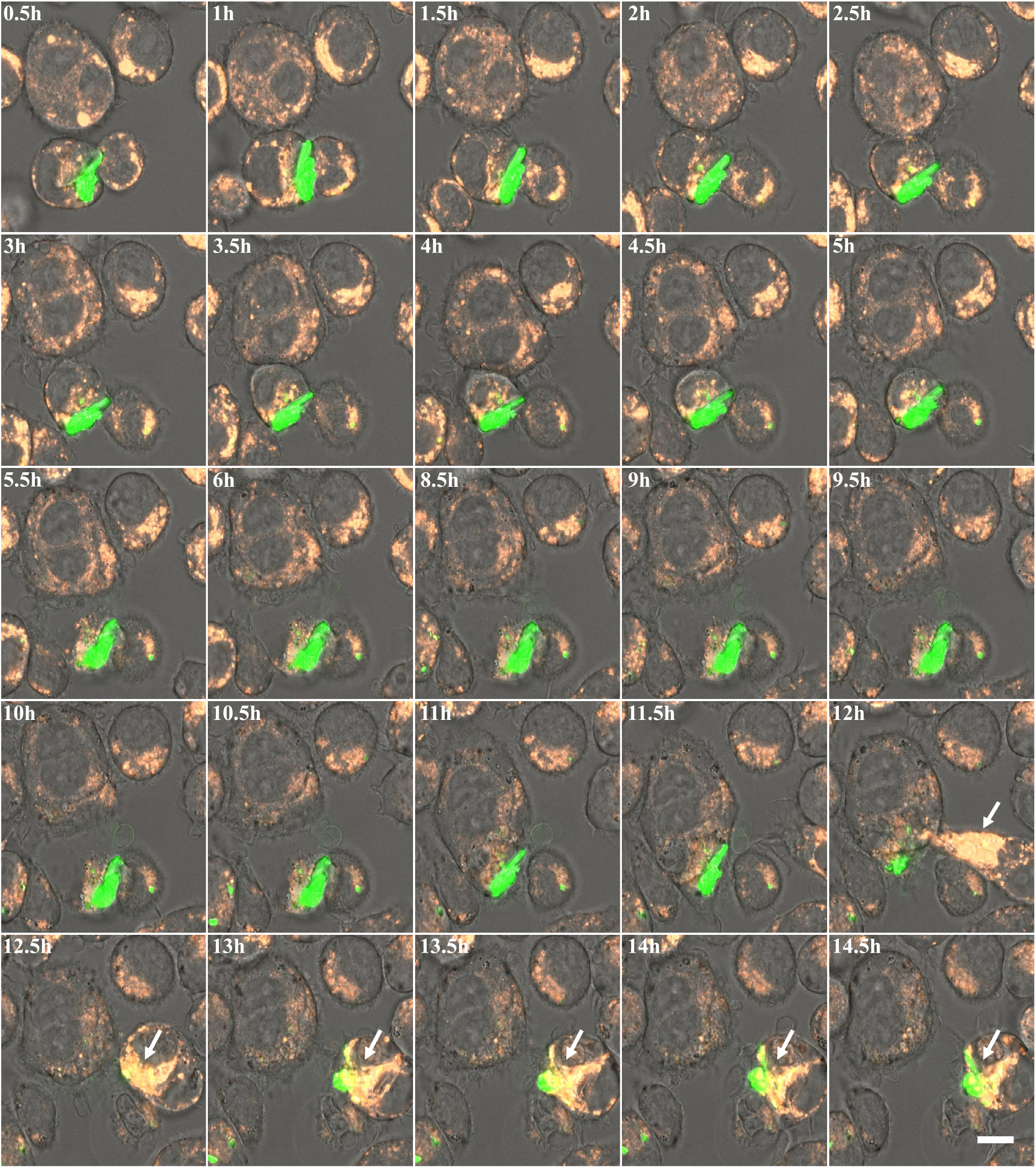
Time-lapse live-cell imaging of the processing of TF CCs by J774 macrophages. Macrophages were incubated overnight with Rh-dextran to label LE/Lys before exposure to TF CCs in phenol red-free DMEM/LPDS medium. Cells were imaged every 30 min for approximately 14.5 h. Representative time points are shown in chronological order, illustrating the dynamic interaction between LE/Lys and TF CCs during crystal processing. Images are shown as overlays of Rh–dextran (orange), TF CCs (green), and brightfield (gray). Scale bar, 10 µm. Crystal size is approximately 13.2 µm × 4.6 µm.

**Figure S3:**
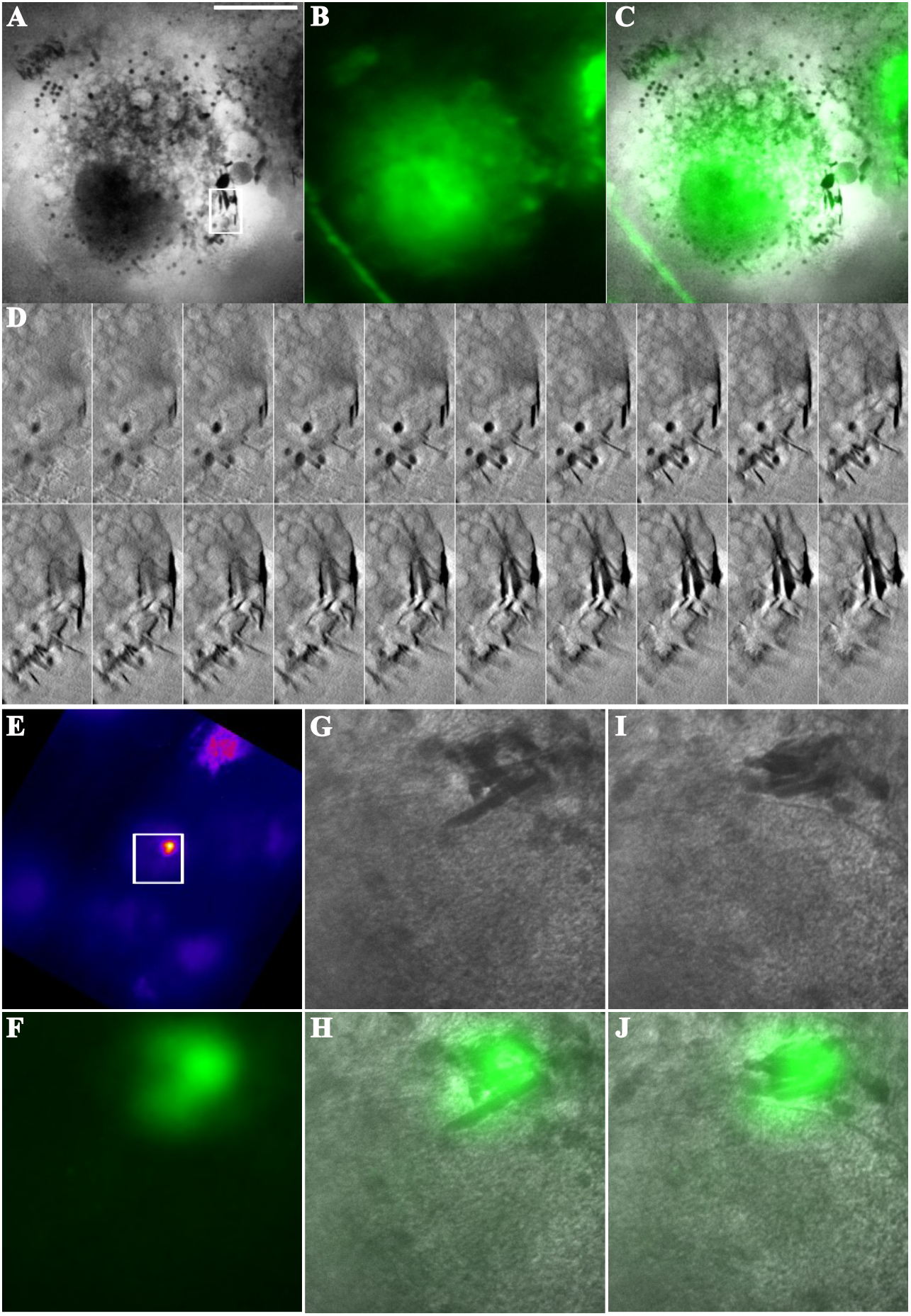
Correlative fluorescence microscopy and Soft X-Ray Tomography (SXT) of TF CCs in macrophages. J774 macrophages were seeded onto grids and cultured for 2 days before overnight incubation with TF CCs. Representative macrophages containing internalized TF CC obtained from Sirius XT, shown as an SXT volume slice (A), epifluorescence image (B), and overlay (C). Scale bar, 10 µm. Higher-magnification views of the crystal in (A-C) are shown in consecutive volume slices (D). SXT images of TF CCs ingested by macrophages were also carried out at the BESSY II synchrotron at Helmholtz Zentrum Berlin (HZB) (E-J). Fluorescence overview image (E), zoom of the square in E (E-J), with the fluorescence image (F) and individual sections along the vertical axis of a 3D reconstruction (G-J) and correlative images of TF-Chol fluorescence in crystals together with the corresponding SXT (G-H, I-J).

**Figure S4:**
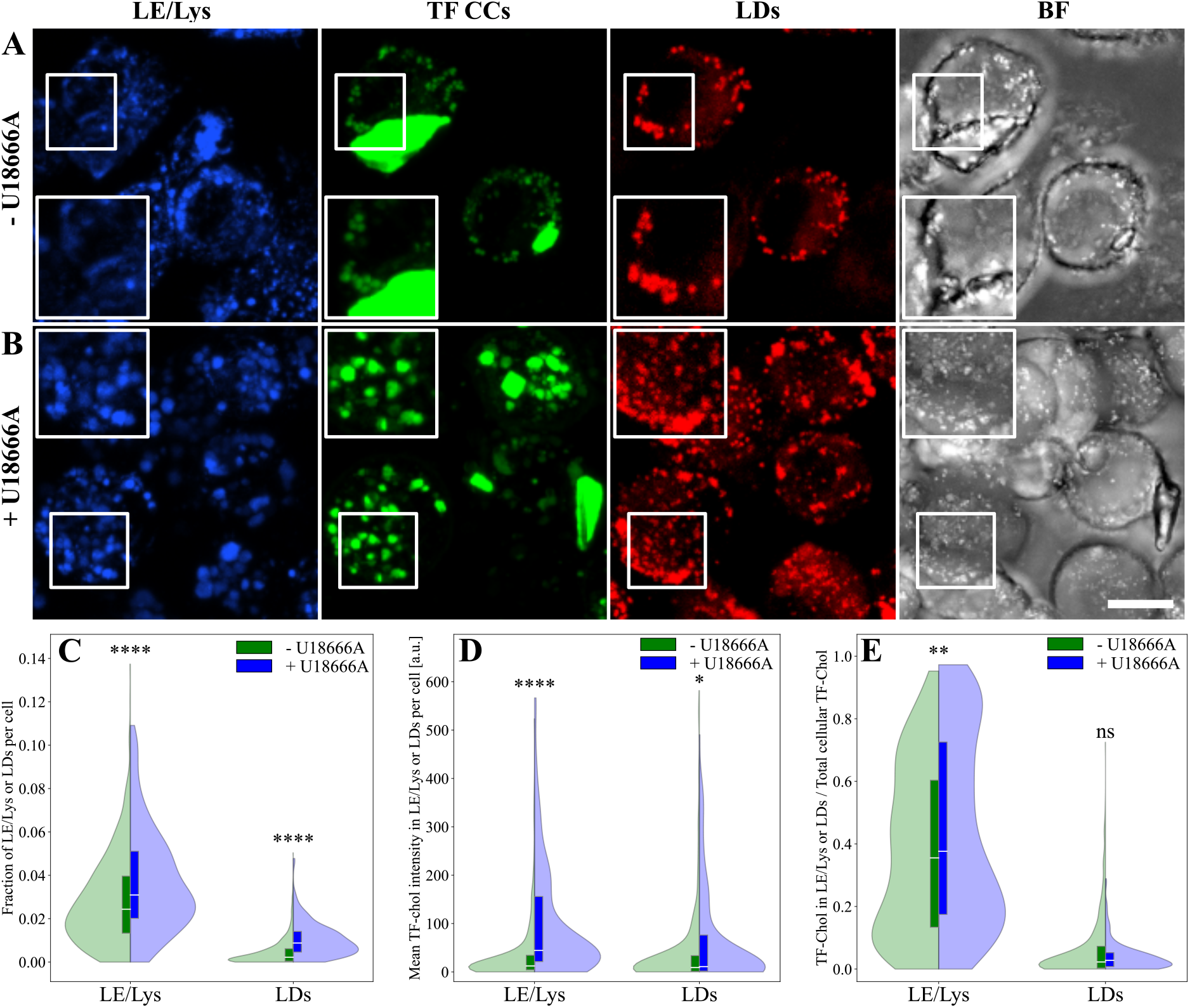
Inhibition of the lysosomal cholesterol transporter NPC1 by U18666A results in TF-Chol accumulation within LE/Lys. J774 macrophages were incubated with TF CCs for 5 h, washed, and either maintained in fresh medium (A) or treated overnight with U18666A (B). Representative confocal maximum intensity projection images are shown. Scale bar, 10 µm. Quantification was performed from five independent experiments for TF CCs-treated cells (n = 488 cells) and three independent experiments for U18666A-treated cells (n = 209 cells). LE/Lys and LD volume fractions, calculated as the number of organelle-positive voxels relative to the total number of voxels within the cell mask, in untreated (green) and U18666A-treated cells (blue) (C). Mean TF-Chol fluorescence intensity in LE/Lys or LDs per cell (D). Fraction of total cellular TF-Chol fluorescence detected within LE/Lys and LDs, calculated as the TF-Chol fluorescence within the respective organelle mask divided by the total TF-Chol fluorescence within the cell (E). Statistical significance was determined using Mann-Whitney U test: p = 6.89*×*10*^−^*^7^ (C, LE/Lys), 5.18*×*10*^−^*^28^ (C, LDs), 9.51*×*10*^−^*^27^ (D, LE/Lys), 1.07 *×* 10*^−^*^2^ (D, LDs), 9.35 *×* 10*^−^*^3^ (E, LE/Lys), and 7.45 *×* 10*^−^*^1^ (E, LDs).

**Figure S5:**
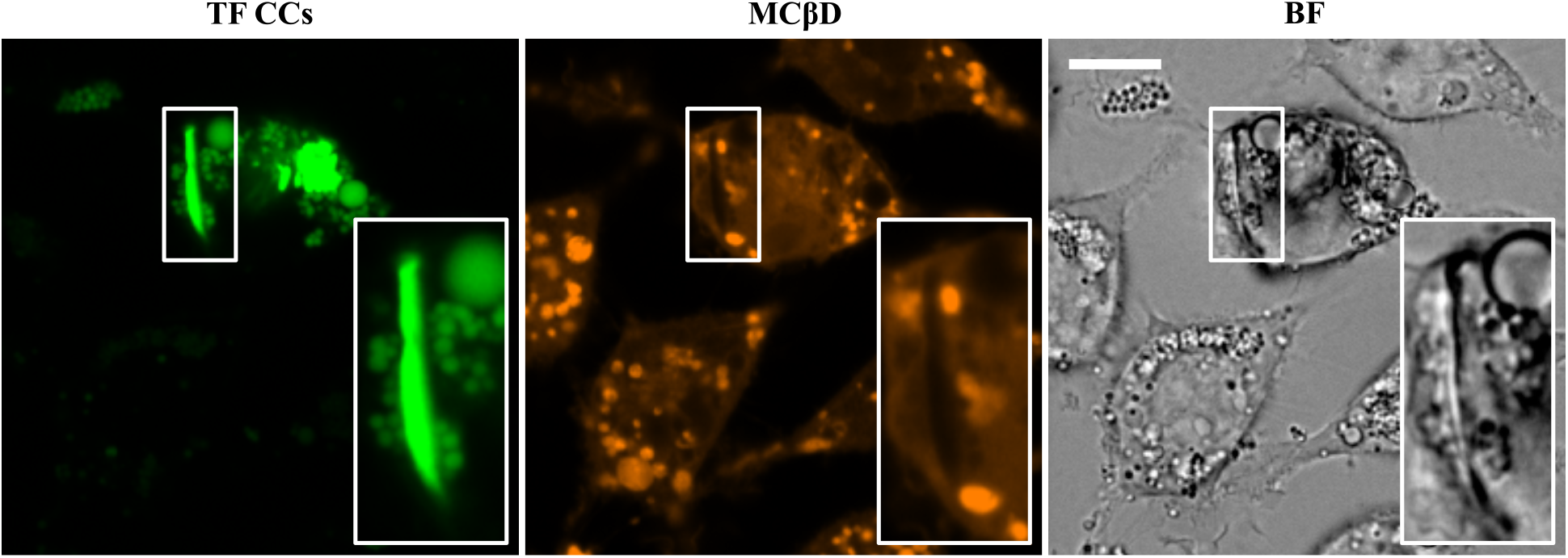
Fluorescent cyclodextrin shows no accumulation at intracellular TF CCs in macrophages. Macrophages were incubated with TF CCs for 5 h to allow crystal uptake, washed, and treated overnight with a mixture of fluorescent and non-fluorescent M*β*CD (1:10, final concentration of 1 mM). Representative fluorescence and bright-field (BF) images are shown. Insets show a magnified view of the boxed region containing a TF CC. Scale bar, 10 µm.

**Figure S6:**
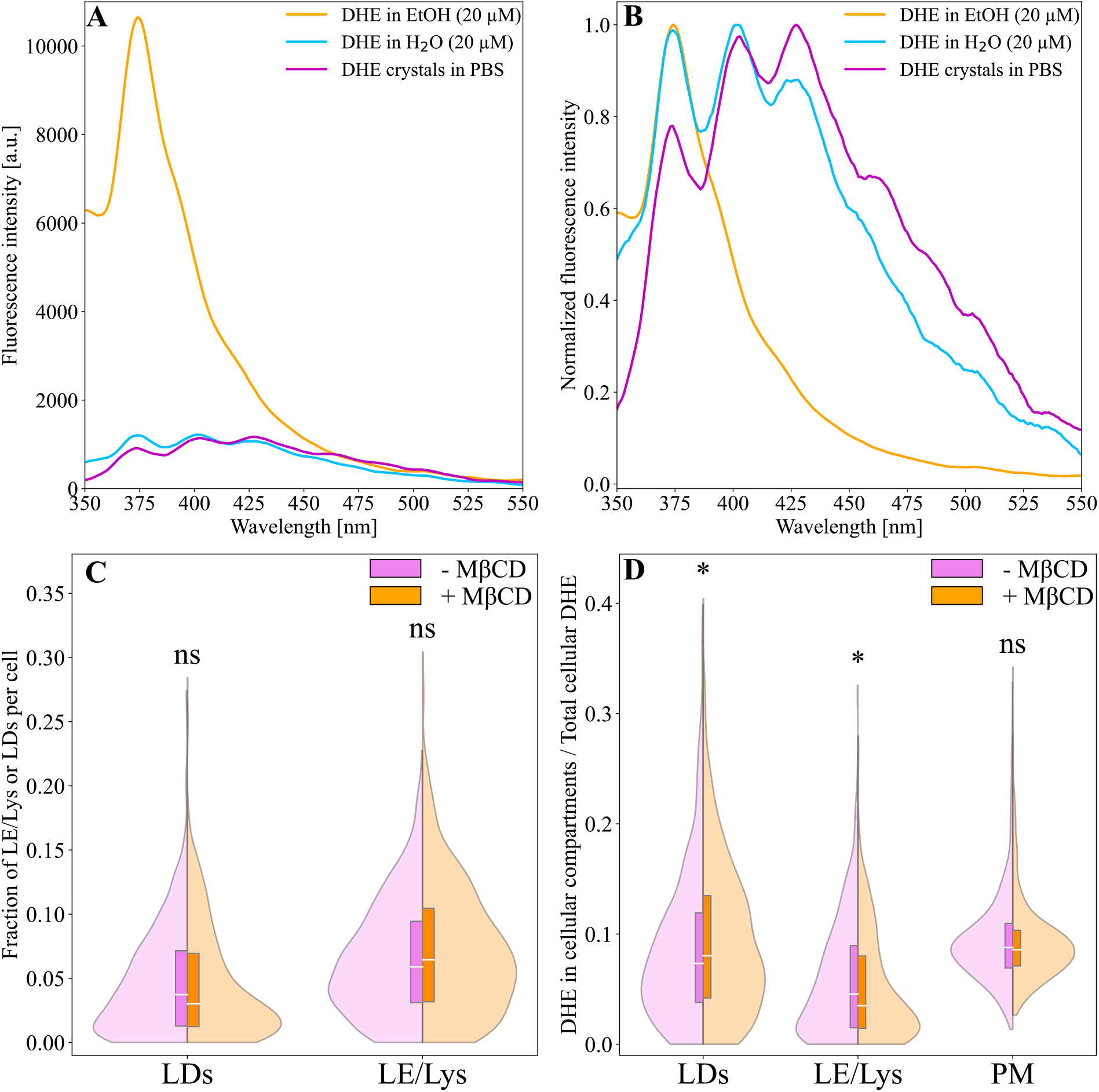
Characterization of DHE crystals and intracellular redistribution of DHE following M*β*CD treatment. Emission spectra of 20 µM DHE in ethanol (EtOH, orange) or water (H_2_O, blue), and DHE crystals (0.08 mg) in PBS (magenta), shown as raw (A) and normalized (B) spectra. For cellular analysis, macrophages were incubated with DHE crystals (0.08 mg) for 5 h, washed, and either maintained overnight in fresh LPDS medium (purple) or treated with M*β*CD (1 mM) (orange). Quantification was performed from three independent experiments (553 cells in the DHE crystal group and 518 cells in the M*β*CD-treated group). The area fraction of LDs (p = 6.08*×*10*^−^*^2^) or LE/Lys (p = 3.4*×*10*^−^*^1^) per cell was quantified (C). The fraction of total cellular DHE fluorescence detected within LDs (p = 3.57 *×* 10*^−^*^2^), LE/Lys (p = 4.46 *×* 10*^−^*^2^) or PM (p = 5.10 *×* 10*^−^*^1^) was calculated as the DHE fluorescence within the respective compartment mask divided by the total DHE fluorescence within the cell (D).

**Figure S7:**
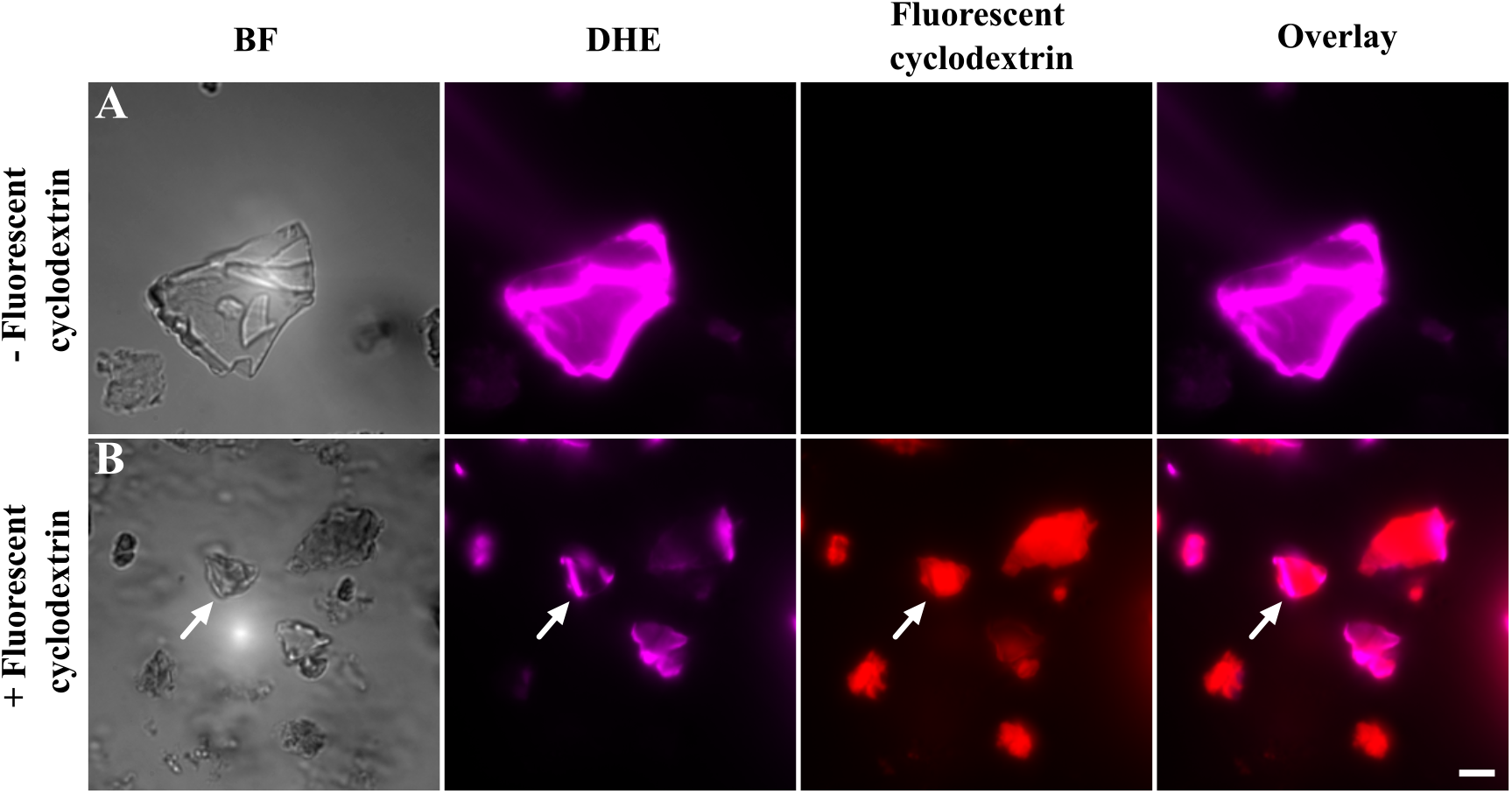
Association of fluorescent cyclodextrin with DHE crystals. Representative UV-widefield images of DHE crystals (0.8 mg) before (A) and after 90 min of incubation with fluorescent cyclodextrin (rhodaminyl-thioureido-RAME*β*) (B). DHE crystals are shown in magenta and fluorescent cyclodextrin in red. White arrows indicate cyclodextrin associated with DHE crystals. Scale bar, 10 µm.

**Figure S8:**
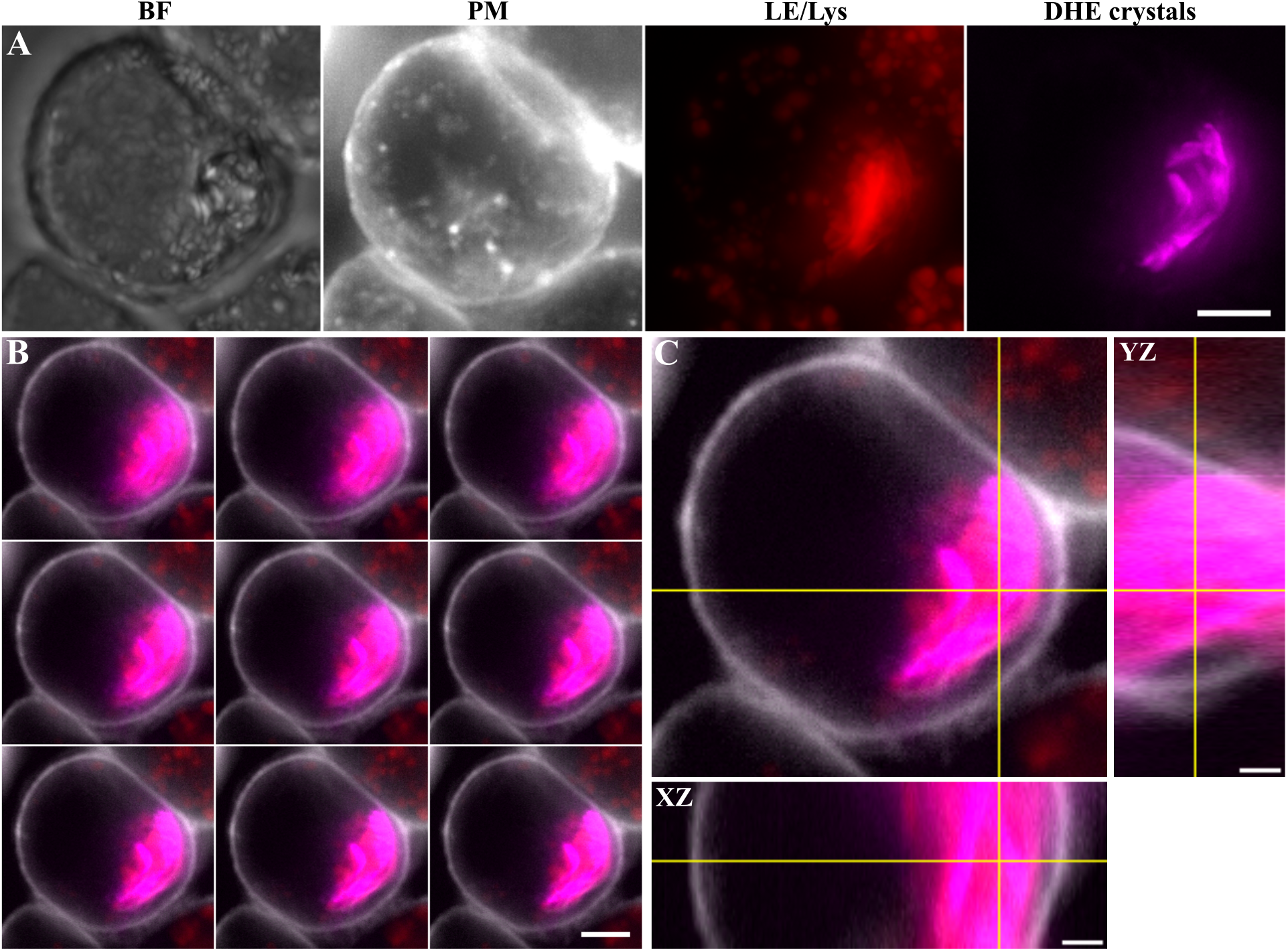
Live-cell 3D imaging of DHE crystals in macrophages. Representative maximum intensity projection images of macrophages incubated overnight with DHE crystals (0.08 mg) (A). Cells were labeled with Rh-dextran (red) to visualize LE/Lys and CellMask (grey) to label the plasma membrane (PM). DHE crystals are shown in magenta. Scale bar, 5 µm. Representative sequential optical sections (Z-stack) through the cell showing the intracellular localization of a DHE crystal. Overlay of the PM (grey), LE/Lys (red), and DHE crystal (magenta). Scale bar, 5 µm (B). Orthogonal views of the selected optical section. Yellow cross indicates the positions of the corresponding XZ and YZ projections shown below and to the right. Scale bars, 2 µm.

**Movie 1. Live-cell 3D imaging of cholesterol crystals in macrophages.** J774 macrophages were incubated overnight with TF CCs (green) and labeled with Rh-dextran (red) to visualize late endosomes/lysosomes (LE/Lys). Sequential optical sections (Z-stack) through the cell demonstrate the intracellular localization of the TF CC and the close association of LE/Lys with the crystal. Scale bar, 5 µm.

**Movie 2. Cyclodextrin promotes dissolution of TF CCs and lipid droplets accumulation in macrophages.** Macrophages were incubated with TF CCs for 5 h to allow crystal uptake, washed, and treated overnight with a mixture of fluorescent and non-fluorescent M*β*CD (1:10, final concentration of 1 mM). The 3D projection illustrates the dissolution of intracellular TF CCs and the accumulation of LDs. LDs are shown in red, and TF-Chol is shown in green.

## Notes

### Competing Interest Statement

The authors have declared no competing interest.

